# Force redistribution in clathrin-mediated endocytosis revealed by coiled-coil force sensors

**DOI:** 10.1101/2021.06.29.450294

**Authors:** Yuan Ren, Jie Yang, Barbara Fujita, Huaizhou Jin, Yongli Zhang, Julien Berro

## Abstract

Forces are central to countless cellular processes, yet *in vivo* force measurement at the molecular scale remains difficult if not impossible. During clathrin-mediated endocytosis, forces produced by the actin cytoskeleton are transmitted to the plasma membrane by a multi-protein coat for membrane deformation. However, the magnitudes of these forces remain unknown. Here, we present new *in vivo* force sensors that induces protein condensation under force. We measured the forces on the fission yeast HIP1R homologue End4p, a protein that links the membrane to the actin cytoskeleton. End4p is under ∼19 pN force near the actin cytoskeleton, ∼ 11 pN near the clathrin lattice, and ∼9 pN near the plasma membrane. Our results demonstrate that forces are collected and redistributed across the endocytic machinery.

**One-Sentence Summary:** New *in vivo* coiled-coil force sensors reveal force redistribution during endocytosis.

## Introduction

Extracellular materials are transported into cells via endocytosis. In eukaryotic cells, clathrin-mediated endocytosis (CME) is the major internalization pathway for nutrients, signaling molecules and pathogenic agents (*1, 2*). It is implicated in numerous diseases including cancer, neurological disorders and virus entry, and is therefore the focus for both basic and translational research (*3*). The distinctive feature of CME is a layer of proteinaceous coat, of which clathrin is a prominent member. The coat contains more than 20 evolutionarily conserved proteins that assemble at the intracellular side of the endocytic site, curves as the endocytic pit matures, and disassembles after the budding of the endocytic vesicle (*4, 5*). Endocytic coat proteins make extensive interactions with each other, with the lipids of the plasma membrane, and with a meshwork of actin filaments that surrounds the endocytic coat (*6*–*9*). Forces produced by actin polymerization are transmitted through adaptor proteins of the endocytic coat to help transform a flat membrane patch into a cargo-filled vesicle into the cytoplasm (*10*–*12*) (Fig. 1a).

**Figure 1.**
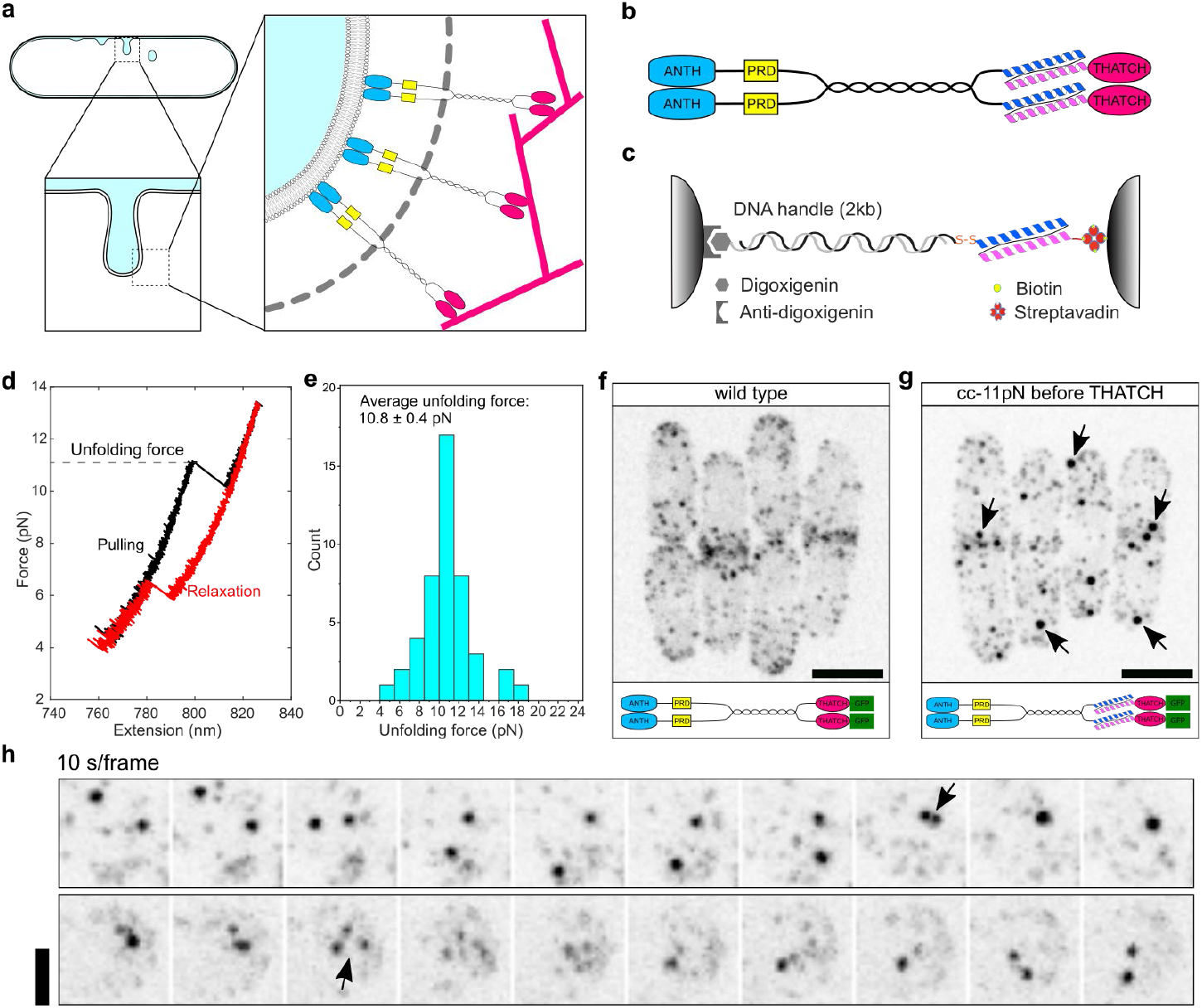
Insertion of a calibrated coiled-coil force sensor within End4p leads to the formation of End4p condensates. **a**, End4p links the lipid membrane to F-actin at sites of clathrin-mediated endocytosis. Upper left, cross section of a fission yeast cell with endocytic pits at different stages. Lower left, side view of an endocytic pit with invaginated membrane. Right, End4p dimers traverses the clathrin coat (gray dotted curve) to transmit the forces generated by the assembly of actin filaments (magenta rods) to deform the plasma membrane. Cyan: End4p lipid binding **A**P180 **N**-**T**erminal **H**omology (ANTH) domain; yellow: End4p proline rich domain; magenta: End4p F-actin binding **t**alin-**H**IP1/R/Sla2p **a**ctin-**t**ethering **C**-terminal **h**omology (THATCH) domain. Drawings are not to scale. **b**, Schematic of an End4p dimer with the insertion of the coiled-coil force sensor before the THATCH domain. **c**, Schematic of the constructs used to calibrate the coiled-coil force sensor with optical tweezers. The coiled-coil is attached to two beads trapped by optical tweezers through a 2260-bp DNA handle. **d**, Representative force-extension curves obtained by pulling (black curve) and then relaxing (red curve) the coiled-coil force sensor. **e**, Distribution of the unfolding force for the coiled-coil force sensor cc-11pN (N=46). The average unfolding force is presented as mean ± SEM. **f**, Representative fission yeast cells expressing fluorescently-tagged wild-type End4p. End4p is mostly found in endocytic patches that are enriched at cell tips and the division plane. See also Fig. S1 and movie 1. **g**, Representative fission yeast cells expressing fluorescently-tagged End4p with cc-11pN inserted before the THATCH domain as in **b**. Multiple large persistent spherical condensates of End4p constructs (indicated by arrows) can be distinguished from transient diffraction limited endocytic patches. **h**, End4p condensates display liquid-like behaviors. Upper panel, fusion of two End4p condensates (arrow). Lower panel, fission of one End4p condensate into two smaller condensates (arrow). See the full dynamics of End4p condensates in Fig. S2, movie 2 and movie 3. **f** and **g**, scale bars: 5 μm. **h**, scale bars: 2 μm. The schematic under each image indicates the modifications on End4p.

Deformation of the membrane during CME is energetically expensive (*12*–*14*). In mammalian cells with elevated membrane tension and in yeast cells where the turgor pressure is as high as 1 MPa, actin polymerization is required for successful CME (*4, 14, 15*). Epsins and Hip1R are believed to transmit the force produced by actin assembly to the plasma membrane because their removal stalls the invagination of endocytic pits, and their C-terminal domains (ACB and THATCH, respectively) bind actin filaments and their N-terminal domains (ENTH and ANTH, respectively) bind PIP2 on the membrane (*8, 10, 11, 16*) (Fig. 1b). Because both proteins also bind clathrin and other endocytic coat proteins, it is possible that forces could be transmitted to the membrane via multiple routes (*6, 17*–*19*). In contrast to the abundant knowledge gleaned from biochemical and genetic approaches, a quantitative understanding of force production and distribution at the molecular level during CME is lacking, because tools to measure forces in live cells in the context of small (∼ 100-nm diameter) and transient (∼10 s) endocytic pits are scarce, difficult to use, and often too bulky to insert into proteins without causing side effects. Here we present a new approach for force measurement *in vivo* that overcomes these limitations.

## Results

The assembly of the endocytic coat relies on weak and multivalent interactions that facilitate the rapid exchange of binding partners during the dynamic rearrangement of the endocytic coat and the constant change in membrane shape during CME (*20*–*22*). Many endocytic proteins contain promiscuous binding sites, multiple copies of short peptide motifs, and intrinsically disordered regions (IDRs) (*9, 20, 21, 23*). Recent developments in protein engineering have indicated that the dual incorporation of IDRs and oligomerization domains (either controlled by light or small molecules) promotes the condensation of proteins *in vivo* (*24*–*26*). We reasoned that if a mechanically actuated oligomerization domain is introduced into an endocytic adaptor protein that contains an IDR, protein condensation could be induced in a force-dependent manner. The force required to activate this domain for oligomerization could be determined through calibration, and the formation of protein condensates would inform us of the presence of force.

Dimeric coiled-coils are excellent candidates for such mechanically actuated oligomerization domain, because they are small, have well-characterized shape and mechanical properties, and can be easily introduced into proteins without affecting their normal functions (*27*–*29*) (Fig. S3a, S4 S8). We and others have previously measured the unfolding forces and energies of dimeric coiled-coils, and found that they unfold at pulling force thresholds in the range of 2-14 piconewtons (pN) (*30*–*32*). The unfolding of a coiled-coil exposes two hydrophobic interaction surfaces that can mediate higher order oligomerization (*33*) (Fig. S3b, e), while a folded coiled-coil puts an upper limit to the local force magnitude and serves as a control for the insertion. Therefore, we used this property to repurpose dimeric coiled-coils as *in vivo* force sensors.

First, we linked the two parallel α-helices in a heterodimeric GCN4 leucine zipper with a 30-amino acid flexible linker to form a single chain polypeptide that can be genetically encoded (Fig. S3a). To measure its unfolding force using optical tweezers, a single GCN4 coiled-coil was tethered between two beads held in two optical traps (Fig. 1c) and pulled to high force by separating the two traps at a speed of 10 nm/s (*31*), which is the typical predicted speed of actin polymerization during endocytosis (see discussion of force vs. speed in the supplementary text). Unfolding of the coiled-coil was manifested by a sudden extension increase at ∼11 pN (Fig. 1d, black trace). The unfolded coiled-coil refolded during relaxation, but at a lower force (Fig. 1d, red trace). Repeated pulling and relaxation reveals a distribution of unfolding forces, with a single peak at 10.8 ± 0.4 pN (Mean ± SEM) (Fig. 1e). This coiled-coil is referred to as cc-11pN hereafter.

We inserted cc-11pN into the fission yeast homolog of Hip1R (End4p) at its genomic locus using CRISPR/Cas9 so that the expression level of End4p was not perturbed (Fig. S4). End4p functions as a dimer *in vivo* (*10, 34*), and contains an IDR between its proline rich domain and its dimerization domain (Fig. S1). In wild-type cells, fluorescently tagged End4p is present at endocytic sites and appear as diffraction-limited puncta (hereafter referred to as End4p patches) that are enriched at cell tips during interphase and around the division plane during mitosis (Fig. 1f, S2a, movie1, S5b-c). When cc-11pN was inserted before the actin-binding THATCH domain of End4p, many large spherical End4p condensates appeared among small patches (Fig. 1g, S5e-f). The condensates remained visible in the cell during the entire time course of the movies (>10 min) whereas End4p remained at endocytic patches for a much shorter time (<50 seconds). Interestingly, the condensates displayed behaviors such as fusion and fission (Fig. 1h, S2b-c, movies 2-3), suggestive of liquid-like features. We hypothesized that condensate assembly of End4p constructs is mediated by force-induced unfolding of cc-11pN, which in turn leads to the formation of inter-molecular association of the unfolded α-helices (Fig. 2a, S3e). To test this hypothesis, we created a construct where we shortened the linker between the two α-helices of cc-11pN to force the coiled-coil to be unfolded (Fig. S3c-d). Indeed, we observed larger condensates (Fig. 2b, S5h-i), supporting our hypothesis that the unfolded coiled coils mediate the condensate assembly. The insertion of cc-11pN did not change the timing and number of End4p molecules recruited to the endocytic sites, and no growth defect was detected, demonstrating that insertion of the coiled-coil force sensors do not disrupt End4p’s function, even when the coiled-coil is open and induces the constructs’ condensation (Fig. S4, S8).

**Figure 2.**
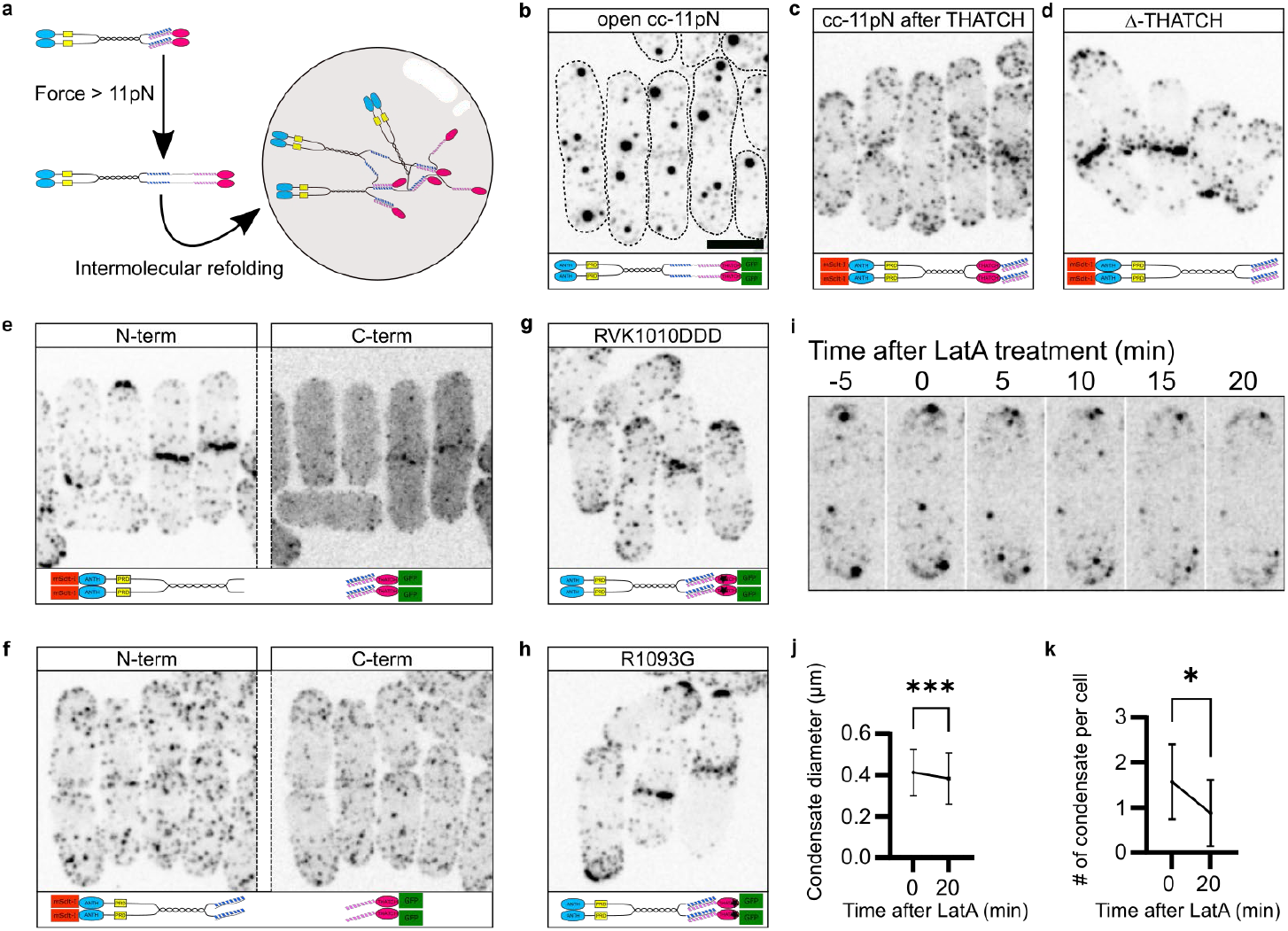
Force-induced unfolding of the coiled-coil force sensor promotes the formation of End4p condensates. **a**, Schematic of End4p condensate formation. The coiled-coil force sensor unfolds when the magnitude of force on End4p exceeds the unfolding force threshold of the coiled-coil force sensor. During the refolding process, α-helices from different End4p molecules mediate the entanglement of End4p molecules into condensates. See also Fig. S3**e. b-i**, Localization of different End4p constructs in fission yeast cells. Both the N- and C-terminal End4p fragments are fluorescently labeled in **e**-**f. b**, The insertion of an open conformation cc-11pN into End4p led to larger End4p condensates. See also Fig. S3**c**-**d**, S5. **c**-**e**, The insertion of cc-11pN into End4p does not lead to condensate formation at the absence of force. The force on cc-11pN was removed by inserting after the THATCH domain (**c**), deleting the THATCH domain (**d**), or disconnecting the THATCH domain with the rest of End4p (**e**). **f**, The formation of End4p condensates is dependent on the connection between the two α-helices of cc-11pN. The cleaving of the linker of cc-11pN prevented the formation of condensates. See also Fig. S3**e**-**f. g**-**h**, End4p does not form condensates when a cc-11pN was inserted before the THATCH domain, and RVK1010DDD (**g**) or R1093G (**h**) mutations were introduced into the THATCH domain to abolish its binding to F-actin. **i**, Snapshots of fission yeast cells with End4p condensates before and after the treatment with 100 μM LatA. The size (**j**) and number(**k**) of End4p condensates decreases after actin assembly is impaired by the latA treatment. Data in **j**-**k** are presented as the mean ± SD (>300 condensates from n = 3 independent repeats). Asterisk indicates a significant difference. **j**, P =0.0003, two-tailed t test. **k**, P =0.01, two-tailed t test. **b-h**, Scale bar in **b** applies to all images, 5 μm. End4p is tagged at the C-terminal with mEGFP in **b**-**i**, and End4p is also tagged at the N-terminal with mScarlet-I in **e**-**f**. *In vivo* protein cleaving in **e**-**f** was achieved by the insertion of a 2A peptide. The schematic under each image indicates the modifications on End4p. See the localization of End4p without cc-11pN in Figure S6.

We constructed several control strains to demonstrate that the formation of End4p condensates is indeed force dependent. To remove the force on cc-11pN, we inserted it at the C-terminal of End4p (Fig. 2c), deleted the THATCH domain (Fig. 2d) or split the construct after the dimerization domain of End4p using the self-cleaving 2A peptide (Fig. 2e). We did not observe any condensate in any of the three cases, suggesting that the incorporation of cc-11pN *per se* does not cause End4p condensation, and that force is needed to generate End4p condensates. Splitting cc-11pN between the two α-helices prohibited the formation of intermolecular coiled-coils, and thereby prevented the formation of End4p condensates (Fig. 2f, S3f). Mutations in the THATCH domain that abolish the binding of End4p to F-actin (RVK1010DDD in Fig. 2g, R1093G in Fig. 2h) also prevented the formation of End4p condensates (*35, 36*). In all these experiments, the cellular localization of these End4p constructs was the same with or without the insertion of cc-11pN (Fig. S6). Last, by inhibiting the polymerization of F-actin and accelerating the disassembly of F-actin through Latrunculin A (LatA)(*37*), we observed a reduction in both the size and the number of End4p condensates (Fig. 2i-k), demonstrating that the formation of End4p condensates depends on F-actin polymerization. Collectively, our data are consistent with the idea that force induced unfolding of cc-11pN in End4p promotes the protein condensation of End4p constructs, and that the magnitude of force on End4p between the dimerization domain and THATCH domain is above 11 pN.

To further assess the force involved in the condensate assembly, we designed and tested four more force sensors that are based on artificial coiled-coils (*38, 39*) and a B-ZIP protein (*40*) (Fig. 3, Fig. S7a) and that have average unfolding force thresholds of 8.2 ± 0.2 pN, 10.0 ± 0.1 pN, 17.5 ± 0.7 pN and 20.0 ± 0.7 pN, which are hereafter referred to as cc-8pN, cc-10pN, cc-18pN and cc-20pN respectively (Fig. 3a, Fig. S7). cc-18pN was derived from cc-20pN by point mutations (Fig. S7a). Insertion of cc-8pN, cc-10pN and cc-18pN into the same position as cc-11pN before the THATCH domain led to the formation of End4p condensates, whereas insertion of cc-20pN did not (Fig. 3b-c). An unfolded version of cc-20pN before the THATCH domain caused End4p condensates (Fig. S7g). Insertion of these four coiled-coils into actin-binding defective End4p did not lead to the formation of End4p condensates (Fig. S7c-f) demonstrating that condensation with cc-8pN, cc-10pN and cc-18pN was force dependent. We also observed a positive correlation between the diameter of End4p condensates and the unfolding force thresholds of the calibrated coiled-coils (Fig. 3b). This correlation is consistent with the idea that the force-induced unfolding of the coiled coils and their subsequent intermolecular association drive condensate formation (Fig. S3). Together, this library of calibrated coiled-coil force sensors showed that the magnitude of force before the THATCH domain is between 18 and 20 pN. Using two published FRET-based *in vivo* force sensors (*34, 41*) we confirmed that force on End4p was larger than 11 pN before the THATCH domain (Fig. 3d-e). Because the FRET-based force sensors measure force up to 11pN, our coiled-coil force sensors had the advantage to achieve force measurements in a range never achieved *in vivo* before.

**Figure 3.**
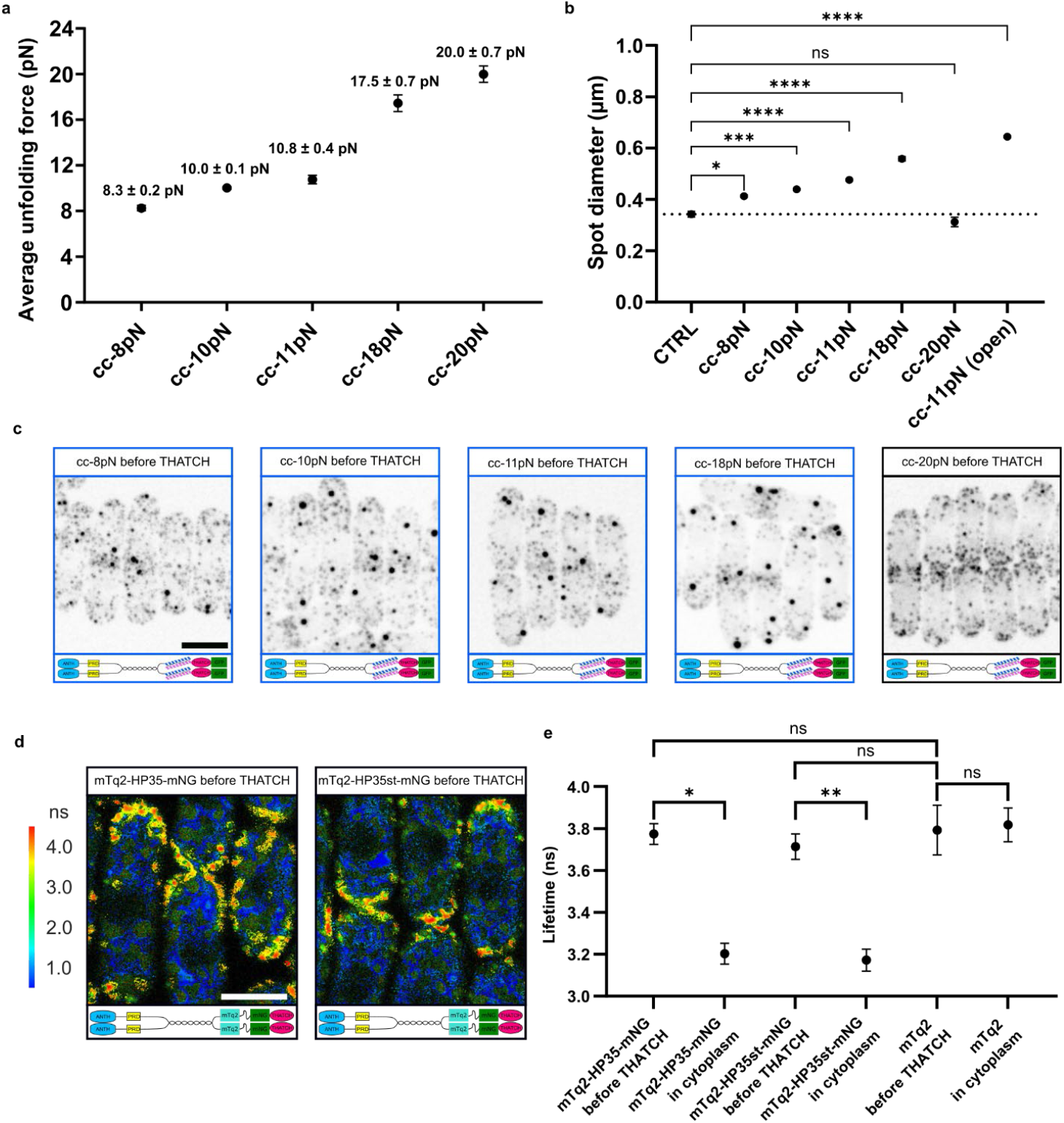
Forces in the 18-20pN range before the Edn4p THATCH domain are measured by a library of calibrated coiled-coil force sensors. **a**, Average unfolding forces of the coiled-coil force sensors, presented as mean ± SEM. N=29 for cc-8pN, N=45 for cc-10pN, N=54 for cc-18pN, N=48 for cc-20pN. See the distribution of unfolding forces and controls of the force sensors in Figure S7. **b**, Quantification of spot diameters in wild type cells and cells with different coiled-coils inserted into End4p before THATCH, presented as mean ± SEM (pooled data from >1000 condensates from at least three independent repeats). Asterisk indicates a significant difference (*: p<0.05, ***: p<0.001, ****: p<0.0001, ordinary one-way ANOVA). The corresponding images of cells can be seen in **c. d**, Donor fluorescent lifetime of fission yeast cells with FRET force sensors inserted into End4p before THATCH. The FRET pair is mTurquoise2 (mTq2, donor) and mNeonGreen (mNG, acceptor). HP and HP35st are peptides that show linear extension to forces in the 6-8 pN and 9-11 pN ranges, respectively(*34*). **e**, Quantification of donor fluorescent lifetime in endocytic patches from cells with the FRET force sensors in End4p before THATCH, donor only in End4p before THATCH, or donor only in the cytoplasm, presented as mean ± SD (pooled data from >50 cells from at least three independent repeats). The donor lifetime of both FRET force sensors in End4p before THATCH is the same as in the donor only control, indicating a lack of FRET and a force magnitude higher than 11pN. Asterisk indicates a significant difference (*: p<0.05, **: p<0.01, Mann-Whitney test). Scale bar in **c** and **d**, 5 μm. The schematic under each image indicates the modifications on End4p.

This library of calibrated coiled-coil force sensors not only allowed us to refine force measurements before the THATCH domain but also to measure the force at two other locations along End4p (Fig. 4a, S1a). The coiled-coil force sensors were inserted after the lipid-binding ANTH domain (Fig. 4a, left column), or between the proline rich domain and the dimerization domain (Fig. 4a, right column). The insertion of a given coiled coil at different locations led to End4p condensates in some but not all cases (compare each row in Fig. 4a). Because the formation of End4p condensates reflects the local presence of force above the unfolding force threshold of the coiled-coil force sensors, our results demonstrate a gradient in the magnitude of force along an End4p molecule: between 8 and 10 pN after the ANTH domain, between 10 and 11 pN between the proline rich domain and the dimerization domain, and between 18 and 20 pN before the THATCH domain (Fig. 4e). These data also provided internal controls for the full functionality of the constructs when the forces are lower than the sensors’ force thresholds, since the timing of endocytosis and the number of End4p molecules were unchanged for all constructs that did not condensate (Fig. S8).

**Figure 4.**
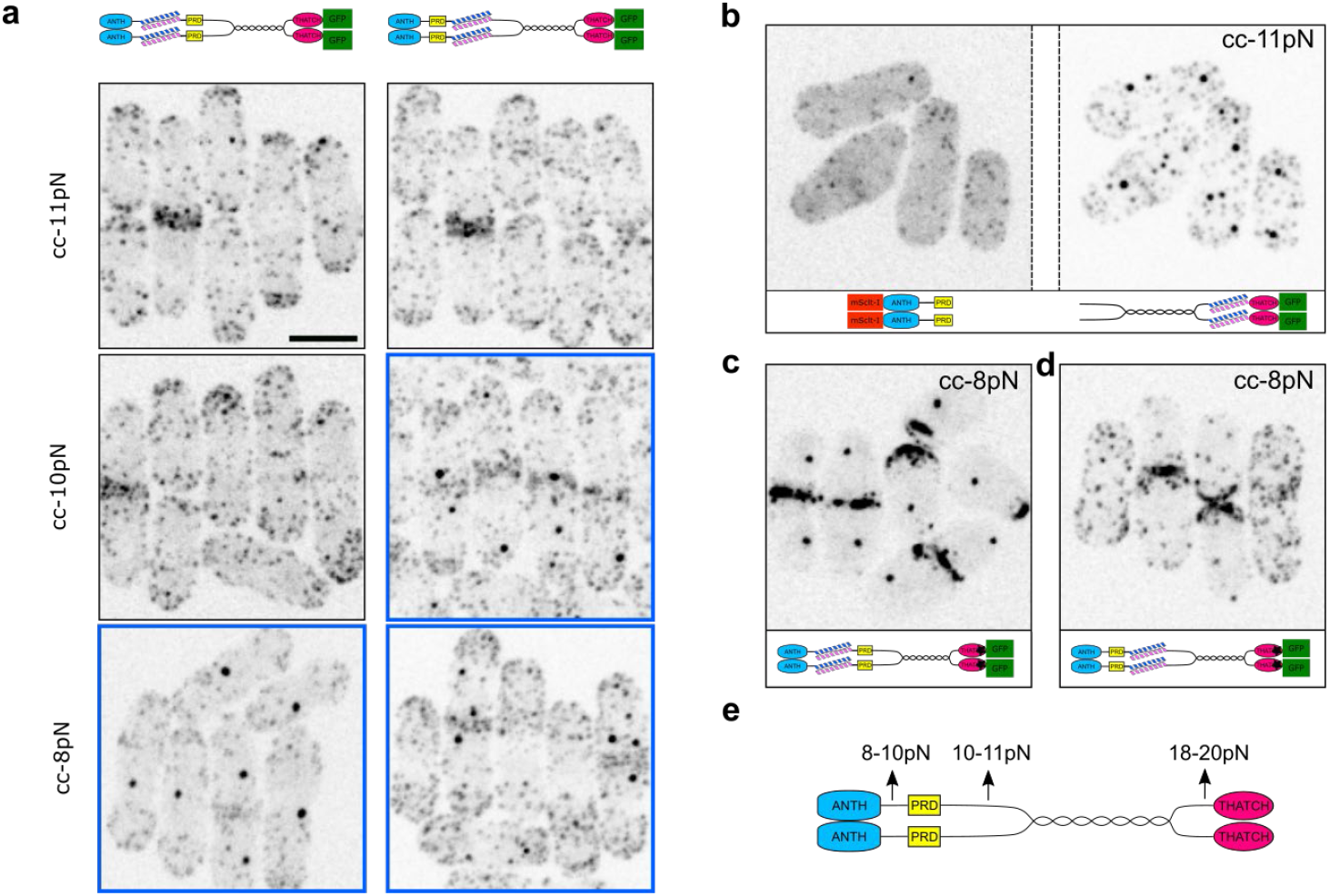
End4p is under a force gradient. **a**, End4p localization when a calibrated coiled-coil force sensor was inserted after the ANTH domain (first column), or after the proline rich domain (second column). Coiled-coils unfolding thresholds are 11 pN (first row), 10 pN (second row), and 8 pN (third row). Images containing End4p condensates are boxed in blue frames. **b**, End4p N- and C-terminal localization when a cc-11pN was inserted before the THATCH domain, and a self-cleaving 2A peptide was introduced after the proline rich domain. The End4p C-terminal fragment formed condensates despite its disconnection from the lipid-binding ANTH domain, demonstrating that at least 11 pN of force is transmitted by End4p C-terminal (after the proline-rich domain) to other components of the endocytic machinery. See the localization of End4p fragments without cc-11pN in Figure S6**c. c**, End4p localization when a cc-8pN was inserted after the ANTH domain, and an R1093G mutation was introduced into the THATCH domain to abolish its binding to actin filaments. This End4p construct formed condensates despite its inability to bind actin, demonstrating that at least 8 pN of force is transmitted by other components of the endocytic machinery to End4p ANTH domain. **d**, End4p localization when a cc-8pN was inserted after the proline rich domain, and an R1093G mutation was introduced into the THATCH domain to abolish its binding to F-actin. This construct didn’t form condensates. This result and the result of panel **c** demonstrate that most of the force not transmitted through the THATCH domain is transmitted through the proline rich domain. **e**, Distribution of forces on End4p. Our calibrated coiled-coil library allowed us to determine that the forces between different domain of End4p are different. **a-d**, Scale bar in **a** applies to all images, 5 μm. End4p is tagged at the C-terminal with mEGFP in **a**-**d**, and End4p is also tagged at the N-terminal with mScarlet-I in **b**. The schematic under each image indicates the modifications on End4p.

The gradient of force on End4p suggests that the binding of other endocytic coat proteins to End4p distributes the force transmitted from F-actin to the lipid membrane. We tested this hypothesis by splitting End4p after its proline rich domain, therefore disconnecting End4p N-terminal from its C-terminal, while preserving the localization of End4p C-terminal at endocytic patches (Fig. S6e). Despite its inability to bind lipids of the clathrin coated pit, forces on End4p C-terminal were still large enough to unfold cc-11pN, confirming that forces are also transmitted by End4p via its binding partners in the endocytic coat (Fig. 4b). Similarly, when we mutated the THATCH domain to prevent the direct binding between End4p and F-actin, we still measured forces higher than 8 pN between End4p’s ANTH and proline rich domains, while forces after the proline rich domain were now smaller than 8 pN (whereas they were larger than 8 pN in wild-type cells) (Fig. 4c, 4d). This result clearly shows that other endocytic coat proteins mediate the transmission of force to the N-terminal of End4p, even when End4p itself is unable to bind F-actin (Fig. 5a). To further demonstrate the extensive inter-connectivity of the endocytic coat, we deleted the F-actin binding domain of another putative force transmitting endocytic protein Ent1p (homologous to Human epsin-1) (*10, 21*), and measured an increase of tension on End4p before the THATCH domain, while forces after the ANTH domain or after the proline rich domain remained unchanged (Fig. 5b). Taken together, these results demonstrate that, contrary to the common belief, forces are not transmitted directly through adaptor proteins such as End4p but are relayed and redistributed by End4p and its binding partners along the different layers of the endocytic coat.

**Figure 5.**
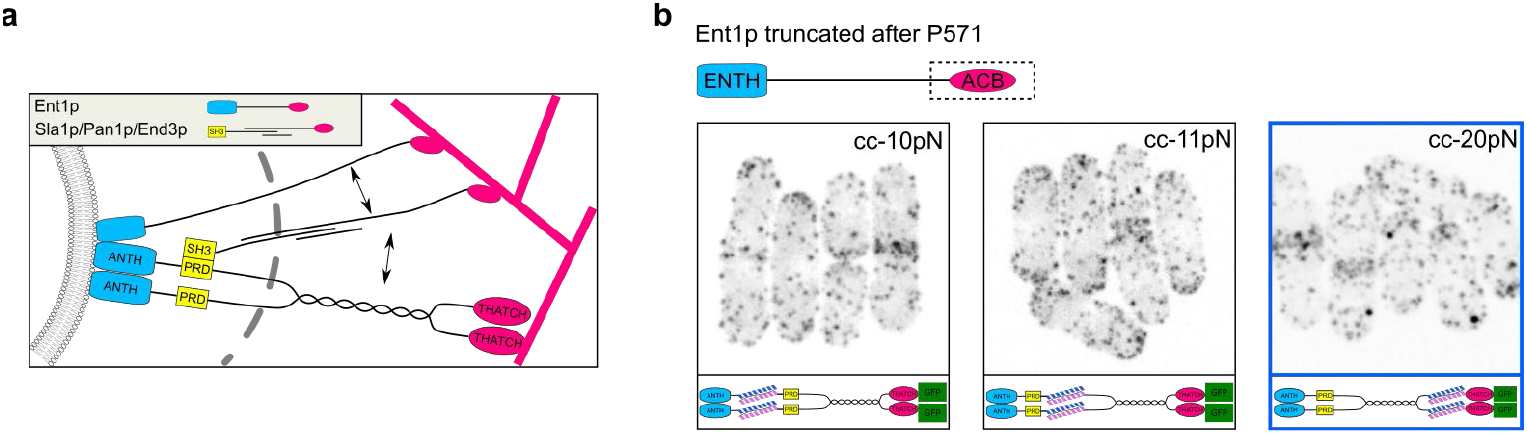
Force redistribution in the endocytic coat is robust against perturbations. **a**, Hypothetical organization of adaptor proteins across the clathrin coat. End4p and Ent1p make direct interactions with the plasma membrane, the clathrin lattice and actin filaments, and are known mechanical linkers to transmit the forces produced by the actin meshwork to deform the plasma membrane. Other endocytic proteins or protein complexes, through their binding to domains in End4p or Ent1p, also relay force from F-actin to the membrane. Double arrows indicate known protein-protein interactions. Drawings are not to scale. **b**, A mutant background was created by removing the actin cytoskeleton-binding (ACB) domain of Ent1p, and calibrated coiled-coil force sensors were inserted after the ANTH domain (first column), after the proline rich domain (second column), or before the THATCH domain (third column). The force before the THATCH domain increased to more than 20pN, whereas forces after the ANTH domain or after the proline rich domain remained unchanged. Scale bar, 5 μm. The image containing End4p condensates are boxed in blue frames. End4p is tagged at the C-terminal with mEGFP.

## Discussion

Force requirement in CME has been the subject of multiple theoretical work, but direct force measurements in the densely woven endocytic coat have been lacking (*12, 42, 43*). Only one recent study in *S. cerevisiae* used FRET-based force sensors to measure the forces on the budding yeast homolog of End4p, Sla2, in a mutant background (*34*). This study measured forces larger than 8 pN near the THATCH actin-binding domain. In our study, we developed calibrated coiled-coils as novel force sensors to induce force-dependent condensation of the endocytic adaptor End4p, and discovered that, in a wild-type background, the forces directly transmitted by the actin network to End4p C-terminal end are between 18 and 20 pN. We also showed that there is a gradient of force along End4p, demonstrating forces are relayed and redistributed across the endocytic coat (Fig 4e).

Our new strategy to measure forces in live cells harnesses the pre-existing multivalent interactions in protein complexes and well complements the current FRET-based approaches (*41, 44*). Our method circumvents the need for inserting two large fluorescent proteins in the middle of the target protein used in the FRET force sensors and provides a larger range of force measurement up to 20 pN *in vivo*. For example, the 18-20 pN force measurement before THATCH had been impossible with the currently existing FRET force sensors (Fig. 3). In addition, the FRET force sensors could not be inserted into two of the three sites within End4p where the coiled-coil force sensors were readily inserted. We suspect End4p is less tolerant for large insertions closer to the plasma membrane where the density of proteins is extremely high due to the hemispherical geometry of the endocytic coat and where interactions with other endocytic coat proteins are numerous (*6, 45*). The self-propagating intermolecular interactions of the coiled-coil force sensors are triggered by transient force on End4p, and the effect of force is amplified both spatially and temporally through the generation of condensates that have micrometer diameters and outlast the lifetime of the endocytic coat. Condensation of End4p does not perturb the endocytic process, because protein condensation likely happens at the end of CME, when the force on End4p has dropped below the coiled-coil refolding threshold, which is markedly smaller than coiled-coil unfolding threshold (Fig. S4, S8). The combination of the small size and the clear readout makes our coiled-coil based strategy a promising tool for easy *in vivo* force measurements in previously inaccessible locations, and an exciting platform for further development. Another advantage of our strategy is that it embeds internal controls and redundancies since all sensors have very similar folds, and every coiled-coil is a sensor as well as a control for other coiled-coils. After insertion into the same position within a protein, a folded coiled-coil force sensor detects the upper limit of the local force magnitude without changing the protein function, while an unfolded coiled-coil force sensor reports the lower limit of force and drives the formation of protein condensates for easy detection (Fig. 3, 4a). Pairwise application of the coiled-coil force sensors demarcates the force range. Nevertheless, our strategy in the current format measures the peak force on a protein and does not allow one to precisely determine the location or the time frame of the force. The spatial and temporal resolution can be added to the calibrated coiled-coils by integration into a FRET pair (*41, 44*), and the measurement of peak force well complements the force measurement from averaged FRET signal (*34*).

We do not expect the coiled-coil force thresholds measured *in vitro* to be notably different *in vivo*, because the mechanical stabilities of the coiled-coils derive mainly from the hydrophobic residues and therefore robust in aqueous environments (Fig. 3) (*28, 31*). In addition, the force-independent effect (e.g. local protein concentration, protein mobility, pH etc.) on protein condensation is controlled by the insertion of different coiled-coils in the same site (Fig. 3b-c; Fig. 4a). For coiled-coils that have hysteresis, we chose the unfolding force instead of the equilibrium force as the threshold for our sensors, because the unfolding of coiled-coils *in vivo* is unlikely in thermal equilibrium due to the long refolding time (Fig. S9, Table S2). The pulling rate only has a small effect on the unfolding force threshold of coiled-coils in the physiological range (see supplementary discussion)(*46*). Because the coiled-coils are either engineered from nuclear proteins or are artificially designed, we do not expect any binding partners in the cytoplasm to compound the force measurement. Lastly, the coiled-coils are inserted into unstructured regions of End4p to avoid domain disruption, and no defect was observed at the protein or cellular level (Fig. S4, S8).

We estimate the peak force on each End4p molecule to be in the 8-10 pN range after the lipid-binding ANTH domain, in the 10-11 pN range between the proline rich domain and the dimerization domain, and in the 18-20 pN range before the THATCH domain (Fig. 3e). The gradient of force along End4p and our mutant data showing that force can still be transmitted to the membrane even if End4p cannot bind actin strongly suggest that the endocytic forces are integrated along the endocytic coat according to a “collect-and-redistribute” mechanism (*6, 41*). Since previous quantitative microscopy studies showed there are up to ∼120 End4p molecules per endocytic coat (*5*), our data suggest that a maximum total force ∼2300 pN is generated by the actin meshwork on End4p at the periphery of the endocytic coat, and ∼1000 pN is transmitted directly to the lipid membrane by End4p ANTH domain, assuming all End4p domains are under force at the same time. These total force estimates on End4p are smaller but in the same order of magnitude as the total forces theory predicted to be required for endocytosis (*13*). Total forces on the plasma membrane could possibly be larger than our estimates since we showed that force transmission in the endocytic coat is relayed through the binding to other adaptor proteins, as fragments of End4p that do not bind actin are still under tension in the endocytic coat (Fig. 4b-c). We speculate that the Sla1p/Pan1p/End3p protein complex, which arrives later than End4p in the assembly of the endocytic coat and has been shown to have interactions with both End4p and Ent1p, bridges the transmission of force from F-actin to End4p (Fig. 5a) (*47, 48*). The clathrin lattice is probably a hub for integrating the transmission of force in the endocytic coat, as End4p, Ent1p, Sla1p and numerous other endocytic adaptor proteins bind to the clathrin lattice, and we detected a change in the magnitude of force on End4p before and after its dimerization domain, which contains the putative clathrin binding site (*18, 19*). The heavily interconnected endocytic coat ensures redundancy to robustly transmit forces deep into the endocytic coat and to the membrane, despite peripheral perturbations closer to the F-actin binding side (Fig. 5b). We expect the magnitudes of forces during CME to be slightly smaller in budding yeast because turgor pressure there is slightly lower, and we expect the forces to be even smaller in mammalian cells where turgor pressure is several orders of magnitude lower. The redundancy in force transmission through the binding of endocytic coat proteins, however, is probably conserved. The “collect and redistribute” mechanism may be a general theme for robust mechanotransduction *in vivo*.

Although End4p condensates only serve as a readout for the coiled-coil force sensors, our *in vivo* data offer a clear mechanism for the condensation of End4p constructs. We show that valency amplification from zero to two following the unfolding of coiled-coils is critical for generating End4p condensates (Fig. 2b, S3, S7g), and that the disconnection of two α-helices, which prevents intermolecular interactions (Fig. 2f), disrupts End4p condensates. Our coiled-coil force sensors therefore represent a new way to induce *in vivo* protein condensates in a force-dependent manner, joining currently existing approaches using light or small molecules (24–26). The prevalence of low-affinity interactions in the endocytic coat and the large force gradient along End4p hint at the potential involvement of force induced protein condensation in normal CME (*49*–*51*), potentiating the formation of End4p condensates after coiled-coil insertion. The exposure of less hydrophobic residues from the unfolding of weaker coiled-coils may be insufficient to drive protein condensation in other systems. The incorporation of IDR and other functional peptides into the coiled-coil backbone could overcome such restraints and enable force induced protein condensation for low abundance or low valency proteins. The ample knowledge related to the design and engineering of coiled-coils pave the road for additional functionalities based on this simple protein motif.

## Materials and Methods

### Protein constructs and purification

The codon optimized DNA of coiled-coils were synthesized (Invitrogen) and cloned into pGEX-6P-1 vectors (Sigma-Aldrich) after PCR amplification and Gibson cloning (New England BioLabs). Constructs were confirmed by DNA sequencing and introduced into BL21 (DE3) competent E. coli cells (New England BioLabs) for protein expression. GST fusion proteins were purified by binding to Glutathione Sepharose 4B beads (GE Healthcare) and the GST tag was removed by cleaving with PreScission Protease (Sigma-Aldrich). The purified proteins were exchanged to the biotinylation buffer containing 25 mM HEPES, 200 mM potassium glutamate with pH 7.7 and biotinylated at the Avi-tag in the presence of 50 μg/mL BirA, 50 mM bicine buffer, pH 8.3, 10 mM ATP, 10 mM magnesium acetate, and 50 μM d-biotin (Avidity) at 4°C overnight.

### Protein-DNA handle crosslinking

The PCR-generated DNA handle used in the single-molecule experiments was 2,260 bp in length and contained a thiol group (-SH) at one end and two digoxigenin moieties at the other end. The DNA handle was crosslinked to the coiled coil protein construct as was described previously (*31*). Briefly, the protein construct was mixed with the DTDP-treated DNA handle in a 50:1 molar ratio in 100 mM phosphate buffer, 500 mM NaCl, pH 8.5 and incubated at room temperature overnight.

### Single-molecule manipulation experiments

All pulling experiments were performed using dual-trap high-resolution optical tweezers as previously described (*31, 32*). Briefly, an aliquot of the crosslinked protein-DNA mixture was mixed with 5 μL 2.1 μm diameter anti-digoxigenin antibody coated polystyrene beads (Spherotech) and incubated at room temperature for 15 min. Then the anti-digoxigenin coated beads and 0.5 μL 1.7 μm diameter streptavidin-coated beads were diluted in 1 mL PBS buffer (137 mM NaCl, 2.7 mM KCl, 8.1 mM Na_2_HPO_4_, 1.8 mM KH_2_PO_4_, pH 7.4) and were separately injected into the top and bottom channels of a homemade microfluidic chamber. Both channels were connected to the central channel with glass tubing. The two beads entering the central channel were caught in two optical traps. The stiffness of each optical tweezer was calibrated by the Brownian motion of the trapped bead and set to ∼0.15 pN/nm by adjusting the power of the trapping laser. After calibration, the two beads were brought close to allow a single protein to be tethered between them. All manipulation experiments were carried out in the PBS buffer supplemented with the oxygen scavenging system (400 mg/mL glucose (Sigma-Aldrich), 0.02 unit/mL glucose oxidase (Sigma-Aldrich), and 0.06 unit/mL catalase). All single molecules were pulled and relaxed by increasing and decreasing, respectively, the trap separation at a speed of 10 nm/s. We chose this speed because during clathrin-mediated endocytosis in yeast ∼100-nm long actin filaments are assembled in ∼10 seconds (*49, 50*), corresponding to an average growth speed of 10nm/s. Since actin dynamics is the main source of force during endocytosis, we expect endocytic proteins participating in force transmission to be under loads moving at ∼10 nm/s. The data was processed by MATLAB codes as were described elsewhere (*31, 32*) and the unfolding forces were determined from the force-extension curves.

### Yeast strains and media

The *S. pombe* strains used in this study are presented in Supplemental Table S1. Strains were constructed using the method described in our previous publication (*51*), and verified by sequencing of the colony PCR products. Fission yeast cells were grown in YE5S (Yeast Extract supplemented with 0.225 g/L of uracil, lysine, histidine, adenine and leucine), and imaged in EMM5S (Edinburgh Minimum media supplemented with 0.225 g/L of uracil, lysine, histidine, adenine and leucine). Yeast cells were grown at 32 °C under 200 rpm shaking overnight to reach exponential phase at OD_595nm_ between 0.3 and 0.5.

### Growth assay

10 μL overnight grown yeast cells were diluted to OD_595nm_ = 0.1 and spotted onto YE5S plates with 10^0^, 10^1^, 10^2^, 10^3^ serial dilutions. Plates were kept at 32-degree or 37-degree incubator for 48 hours before imaging.

### Microscopy

Cells were imaged on 25% gelatin pad at room temperature on a Nikon TiE inverted microscope (Nikon, Tokyo, Japan) with a CSU-W1 Confocal Scanning Unit (Yokogawa Electric Corporation, Tokyo, Japan) under a CFI Plan Apo 100X/1.45NA Phase objective (Nikon, Tokyo, Japan). Images were acquired with an iXon Ultra888 EMCCD camera (Andor, Belfast, UK). mEGFP tagged strains were excited with a 488-nm argon-ion laser and filtered by Spectra X with a single band pass filter 510/25. mScarlet-I tagged strains were excited with a 561-nm argon-ion laser and filtered by Spectra X with a single band pass filter 575/25. Fluorescent signals from the whole cell were collected with 21 optical sections separated by 0.5 μm, and max projected to create 2D images. The laser excitation and image acquisition settings for the same fluorophore (mEGFP or mScarlet-I) were the same for all strains imaged. Images were displayed and analyzed with the Fiji distribution of ImageJ (NIH, USA).

### Fluorescent lifetime imaging

Cells containing a FRET force sensor (mTq2-HP35-mNG or mTq2-HP35st-mNG, gift from Dr. Michal Skruzny) or the donor only controls were imaged on 25% gelatin pad at room temperature on a Stellaris 8 Falcon laser scanning microscope (Leica, Germany) equipped with a 100X objective with 20% 440nm laser excitation at 600 speed, 512X512 pixels, 1AU (0.896 μm section), 4X line averaging and bidirectional scan. Photons were detected by Leica HyD2 detector. Fluorescent lifetime was calculated with the FLIM module in Leica Application Suite (LAS) X.

### LatA treatment

Cells were washed with EMM5S and loaded into CellASIC microfluidics chambers (Y04C-02-5PK, Millipore-Sigma, Saint Louis, USA) before LatA (Thermo Fisher, MA, USA) treatment. The media exchange was controlled by the CellASIC ONIX2 microfluidics system (Millipore-Sigma, Saint Louis, USA) with a flow pressure of 4 psi. LatA was diluted in EMM5S to a final concentration of 100 μM. Fluorescent signals from the whole cell were collected with 21 consecutive optical sections separated with 0.5 μm z-steps, and stacks were displayed using average intensity projection. Cells were imaged every 5 minutes, and the final images were corrected for photobleaching (exponential fit) before being analyzed.

### Protein condensate analysis

For the quantification of End4p protein condensates, we created an ImageJ plugin that identifies protein condensates and excludes End4p patches based on fluorescence intensity (900-65535), size (4-500 pixel) and circularity (0.2-1.0). Our plugin shows good agreement with manual selection. Examples of automatically identified protein condensates are shown in Figure S5.

### Patch tracking

Following EMM5S washes, cells were loaded into the chambers of a six-well iBidi μslide (iBidi, Munich, Germany) pre-treated with 0.1% poly-l-lysine (Peptide Instituted, Osaka, Japan) for imaging. The fluorescence signal from 5 consecutive optical sections, separated by 0.5 μm z-steps and centered at the mid-plane of the cells, was acquired at one second intervals for one minute. Patch tracking was performed as described previously in (*55, 56*). In brief, after corrections for uneven field illumination and camera noise, the temporal evolution of fluorescence intensity was tracked and measured in the z-sum projected movies with an updated version of the PatchTrackingTools (*55, 56*) for the Fiji distribution of ImageJ (NIH, USA)(*57*). Tracks from each strain are obtained from several movies, aligned, and averaged using temporal super-resolution alignment (*55*) with Matlab R2019a (Mathworks, Natick, USA). We used a calibration curve to convert fluorescence intensity into number of molecules (*55*). The curves depicting the temporal evolution of protein copy number were generated in Matlab and the statistical comparison between strains was performed using Welch’s t-test on the mean peak number of molecules as shown in the accompanying table in Fig. S8.

## Acknowledgments

We thank the Yale West Campus Imaging Core for providing access to the microscopes, and Keck DNA Sequencing Facility at Yale for their assistance. We thank Michal Skruzny for sharing *S. cerevisiae* strains. We thank Jim Rothman, Martin Schwartz, Min Wu, Xiaolei Su, Mathis Riehle, Daniel Suter, Yue Liu, Valentina Greco, Tom Pollard, and Avinash Kumar for their comments on the manuscript.

## Author contributions

Conceptualization: JB, YR

Methodology: YR, JB, YZ

Investigation: YR, JY, HJ

Visualization: YR, JB, JY, YZ, BF

Funding acquisition: JB, YZ

Project administration: JB, YZ

Supervision: JB, YZ

Writing – original draft: YR

Writing – review & editing: YR, JB, JY, YZ

## Data and materials availability

All data needed to evaluate the conclusions in the paper are present in the paper and/or the Supplementary Materials. *S. pombe* strains, and plasmids containing the coiled-coil force sensors are available from J.B. upon reasonable request, and purified protein constructs for single-molecule force calibration are available from Y.Z. upon reasonable request.

## Supplementary Materials

Supplemental text

Figs. S1 to S10

Table S1 to S2

Captions for Movies S1 to S3

## References

1. M. Mettlen, P.-H. Chen, S. Srinivasan, G. Danuser, S. L. Schmid, Regulation of Clathrin-Mediated Endocytosis. Annual Review of Biochemistry. 87, 871–896 (2018).

2. M. Kaksonen, A. Roux, Mechanisms of clathrin-mediated endocytosis. Nat Rev Mol Cell Biol. 19, 313–326 (2018).

3. H. T. McMahon, E. Boucrot, Molecular mechanism and physiological functions of clathrin-mediated endocytosis. Nat Rev Mol Cell Biol. 12, 517–533 (2011).

4. W. Kukulski, M. Schorb, M. Kaksonen, J. A. G. Briggs, Plasma Membrane Reshaping during Endocytosis Is Revealed by Time-Resolved Electron Tomography. Cell. 150, 508–520 (2012).

5. Y. Sun, J. Schöneberg, X. Chen, T. Jiang, C. Kaplan, K. Xu, T. D. Pollard, D. G. Drubin, Direct comparison of clathrin-mediated endocytosis in budding and fission yeast reveals conserved and evolvable features. eLife. 8, e50749 (2019).

6. M. Skruzny, E. Pohl, S. Gnoth, G. Malengo, V. Sourjik, The protein architecture of the endocytic coat analyzed by FRET microscopy. Molecular Systems Biology. 16, e9009 (2020).

7. V. Legendre-Guillemin, ENTH/ANTH proteins and clathrin-mediated membrane budding. Journal of Cell Science. 117, 9–18 (2004).

8. J. J. Baggett, K. E. D’Aquino, B. Wendland, The Sla2p Talin Domain Plays a Role in Endocytosis in Saccharomyces cerevisiae. Genetics. 165, 1661–1674 (2003).

9. L. Maldonado-Báez, B. Wendland, Endocytic adaptors: recruiters, coordinators and regulators. Trends in Cell Biology. 16, 505–513 (2006).

10. M. Skruzny, T. Brach, R. Ciuffa, S. Rybina, M. Wachsmuth, M. Kaksonen, Molecular basis for coupling the plasma membrane to the actin cytoskeleton during clathrin-mediated endocytosis. Proc Natl Acad Sci U S A. 109, E2533–E2542 (2012).

11. M. Skruzny, A. Desfosses, S. Prinz, S. O. Dodonova, A. Gieras, C. Uetrecht, A. J. Jakobi, M. Abella, W. J. H. Hagen, J. Schulz, R. Meijers, V. Rybin, J. A. G. Briggs, C. Sachse, M. Kaksonen, An Organized Co-assembly of Clathrin Adaptors Is Essential for Endocytosis. Developmental Cell. 33, 150–162 (2015).

12. M. Lacy, R. Ma, N. Ravindra, B.-J. letters, Molecular mechanisms of force production in clathrin-mediated endocytosis. FEBS letters (2018).

13. R. Ma, J. Berro, Endocytosis against high turgor pressure is made easier by partial coating and freely rotating base. Biophys J. 120, 1625–1640 (2021).

14. R. Basu, E. L. Munteanu, F. Chang, Role of turgor pressure in endocytosis in fission yeast. Mol Biol Cell. 25, 679–687 (2014).

15. S. Aghamohammadzadeh, K. R. Ayscough, Differential requirements for actin during yeast and mammalian endocytosis. Nat Cell Biol. 11, 1039–1042 (2009).

16. M. Messa, R. Fernández-Busnadiego, E. Sun, H. Chen, H. Czapla, K. Wrasman, Y. Wu, G. Ko, T. Ross, B. Wendland, P. De Camilli, Epsin deficiency impairs endocytosis by stalling the actin-dependent invagination of endocytic clathrin-coated pits. eLife. 3 (2013), doi:10.7554/eLife.03311.

17. R. C. Aguilar, H. A. Watson, B. Wendland, The Yeast Epsin Ent1 Is Recruited to Membranes through Multiple Independent Interactions. J. Biol. Chem. 278, 10737–10743 (2003).

18. Å. E. Y. Engqvist-Goldstein, R. A. Warren, M. M. Kessels, J. H. Keen, J. Heuser, D. G. Drubin, The actin-binding protein Hip1R associates with clathrin during early stages of endocytosis and promotes clathrin assembly in vitro. J Cell Biol. 154, 1209–1224 (2001).

19. J. D. Wilbur, C.-Y. Chen, V. Manalo, P. K. Hwang, R. J. Fletterick, F. M. Brodsky, Actin Binding by Hip1 (Huntingtin-interacting Protein 1) and Hip1R (Hip1-related Protein) Is Regulated by Clathrin Light Chain. J Biol Chem. 283, 32870–32879 (2008).

20. S. M. Smith, M. Baker, M. Halebian, C. J. Smith, Weak Molecular Interactions in Clathrin-Mediated Endocytosis. Front. Mol. Biosci. 4 (2017), doi:10.3389/fmolb.2017.00072.

21. M. M. Garcia-Alai, J. Heidemann, M. Skruzny, A. Gieras, H. D. T. Mertens, D. I. Svergun, M. Kaksonen, C. Uetrecht, R. Meijers, Epsin and Sla2 form assemblies through phospholipid interfaces. Nature Communications. 9, 328 (2018).

22. Y. Zhuo, K. E. Cano, L. Wang, U. Ilangovan, A. P. Hinck, R. Sousa, E. M. Lafer, Nuclear Magnetic Resonance Structural Mapping Reveals Promiscuous Interactions between Clathrin-Box Motif Sequences and the N-Terminal Domain of the Clathrin Heavy Chain. Biochemistry. 54, 2571–2580 (2015).

23. Y. Miao, T. Tipakornsaowapak, L. Zheng, Y. Mu, E. Lewellyn, Phospho-regulation of intrinsically disordered proteins for actin assembly and endocytosis. The FEBS Journal. 285, 2762–2784 (2018).

24. D. Bracha, M. T. Walls, C. P. Brangwynne, Probing and engineering liquid-phase organelles. Nature Biotechnology. 37, 1435–1445 (2019).

25. D. Bracha, M. T. Walls, M.-T. Wei, L. Zhu, M. Kurian, J. L. Avalos, J. E. Toettcher, C. P. Brangwynne, Mapping Local and Global Liquid Phase Behavior in Living Cells Using Photo-Oligomerizable Seeds. Cell. 175, 1467–1480.e13 (2018).

26. H. Nakamura, A. A. Lee, A. S. Afshar, S. Watanabe, E. Rho, S. Razavi, A. Suarez, Y.-C. Lin, M. Tanigawa, B. Huang, R. DeRose, D. Bobb, W. Hong, S. B. Gabelli, J. Goutsias, T. Inoue, Intracellular production of hydrogels and synthetic RNA granules by multivalent molecular interactions. Nat Mater. 17, 79–89 (2018).

27. M. E. Tanenbaum, L. A. Gilbert, L. S. Qi, J. S. Weissman, R. D. Vale, A protein tagging system for signal amplification in gene expression and fluorescence imaging. Cell. 159, 635–646 (2014).

28. L. Truebestein, T. A. Leonard, Coiled-coils: The long and short of it. Bioessays. 38, 903–916 (2016).

29. T. Lebar, D. Lainšček, E. Merljak, J. Aupič, R. Jerala, A tunable orthogonal coiled-coil interaction toolbox for engineering mammalian cells. Nat Chem Biol. 16, 513–519 (2020).

30. M. Goktas, C. Luo, R. M. A. Sullan, A. E. Bergues-Pupo, R. Lipowsky, A. V. Verde, K. G. Blank, Molecular mechanics of coiled coils loaded in the shear geometry. Chem. Sci. 9, 4610–4621 (2018).

31. Y. Gao, G. Sirinakis, Y. Zhang, Highly Anisotropic Stability and Folding Kinetics of a Single Coiled Coil Protein under Mechanical Tension. J. Am. Chem. Soc. 133, 12749–12757 (2011).

32. Z. Xi, Y. Gao, G. Sirinakis, H. Guo, Y. Zhang,Single-molecule observation of helix staggering, sliding, and coiled coil misfolding. PNAS. 109, 5711–5716 (2012).

33. M. J. Pandya, G. M. Spooner, M. Sunde, J. R. Thorpe, A. Rodger, D. N. Woolfson, Sticky-End Assembly of a Designed Peptide Fiber Provides Insight into Protein Fibrillogenesis. Biochemistry. 39, 8728–8734 (2000).

34. M. Abella, L. Andruck, G. Malengo, M. Skruzny, Actin-generated force applied during endocytosis measured by Sla2-based FRET tension sensors. Developmental Cell. 56, 2419–2426.e4 (2021).

35. R. O. McCann, S. W. Craig, The I/LWEQ module: a conserved sequence that signifies Factin binding in functionally diverse proteins from yeast to mammals. Proc Natl Acad Sci U S A. 94, 5679–5684 (1997).

36. T. J. Brett, V. Legendre-Guillemin, P. S. McPherson, D. H. Fremont, Structural definition of the F-actin–binding THATCH domain from HIP1R. Nat Struct Mol Biol. 13, 121–130 (2006).

37. I. Fujiwara, M. E. Zweifel, N. Courtemanche, T. D. Pollard, Latrunculin A Accelerates Actin Filament Depolymerization in Addition to Sequestering Actin Monomers. Curr Biol. 28, 3183–3192.e2 (2018).

38. E. K. O’Shea, K. J. Lumb, P. S. Kim, Peptide ‘Velcro’: Design of a heterodimeric coiled coil. Current Biology. 3, 658–667 (1993).

39. D. G. Gurnon, J. A. Whitaker, M. G. Oakley, Design and Characterization of a Homodimeric Antiparallel Coiled Coil. J. Am. Chem. Soc. 125, 7518–7519 (2003).

40. J. R. Moll, S. B. Ruvinov, I. Pastan, C. Vinson, Designed heterodimerizing leucine zippers with a ranger of pIs and stabilities up to 10− 15 M. Protein Sci. 10, 649–655 (2001).

41. P. Ringer, A. Weißl, A.-L. Cost, A. Freikamp, B. Sabass, A. Mehlich, M. Tramier, M. Rief, C. Grashoff, Multiplexing molecular tension sensors reveals piconewton force gradient across talin-1. Nature Methods. 14, 1090–1096 (2017).

42. S. Dmitrieff, F. Nédélec, Membrane Mechanics of Endocytosis in Cells with Turgor. PLOS Computational Biology. 11 (2015), doi:10.1371/journal.pcbi.1004538.

43. M. Nickaeen, J. Berro, T. D. Pollard, B. M. Slepchenko, Simulation of the mechanics of actin assembly during endocytosis in yeast. bioRxiv, 518423 (2019).

44. C. Grashoff, B. D. Hoffman, M. D. Brenner, R. Zhou, M. Parsons, M. T. Yang, M. A. McLean, S. G. Sligar, C. S. Chen, T. Ha, M. A. Schwartz, Measuring mechanical tension across vinculin reveals regulation of focal adhesion dynamics. Nature. 466, 263 (2010).

45. M. Mund, J. A. van der Beek, J. Deschamps, S. Dmitrieff, P. Hoess, J. L. Monster, A. Picco, F. Nédélec, M. Kaksonen, J. Ries, Systematic Nanoscale Analysis of Endocytosis Links Efficient Vesicle Formation to Patterned Actin Nucleation. Cell. 174, 884–896.e17 (2018).

46. X. Wang, T. Ha, Defining single molecular forces required to activate integrin and notch signaling. Science. 340, 991–994 (2013).

47. Y. Sun, N. Leong, T. Wong, D. Drubin, A Pan1/End3/Sla1 complex links Arp2/3-mediated actin assembly to sites of clathrin-mediated endocytosis. Molecular Biology of the Cell. 26, 3841–3856 (2015).

48. H.-Y. Tang, J. Xu, M. Cai, Pan1p, End3p, and Sla1p, Three Yeast Proteins Required for Normal Cortical Actin Cytoskeleton Organization, Associate with Each Other and Play Essential Roles in Cell Wall Morphogenesis. Molecular and Cellular Biology. 20, 12–25 (2000).

49. M. Kozak, M. Kaksonen, “Phase separation of Ede1 promotes the initiation of endocytic events” (preprint, Cell Biology, 2019),, doi:10.1101/861203.

50. K. J. Day, G. Kago, L. Wang, J. B. Richter, C. C. Hayden, E. M. Lafer, J. C. Stachowiak, Liquid-like protein interactions catalyse assembly of endocytic vesicles. Nat Cell Biol. 23, 366–376 (2021).

51. L.-P. Bergeron-Sandoval, S. Kumar, H. K. Heris, C. Chang, C. E. Cornell, S. L. Keller, P. François, A. G. Hendricks, A. J. Ehrlicher, R. V. Pappu, S. W. Michnick, Proteins with prion-like domains can form viscoelastic condensates that enable membrane remodeling and endocytosis. bioRxiv, 145664 (2021).

52. V. Sirotkin, J. Berro, K. Macmillan, L. Zhao, T. D. Pollard, Quantitative analysis of the mechanism of endocytic actin patch assembly and disassembly in fission yeast. Mol Biol Cell. 21, 2894–2904 (2010).

53. J. Berro, V. Sirotkin, T. D. Pollard, Mathematical Modeling of Endocytic Actin Patch Kinetics in Fission Yeast: Disassembly Requires Release of Actin Filament Fragments. MBoC. 21, 2905–2915 (2010).

54. R. Fernandez, J. Berro, Use of a fluoride channel as a new selection marker for fission yeast plasmids and application to fast genome editing with CRISPR/Cas9. Yeast. 33, 549–557 (2016).

55. J. Berro, T. D. Pollard, Local and global analysis of endocytic patch dynamics in fission yeast using a new “temporal superresolution” realignment method. Mol Biol Cell. 25, 3501–3514 (2014).

56. J. Lemière, Y. Ren, J. Berro, Rapid adaptation of endocytosis, exocytosis, and eisosomes after an acute increase in membrane tension in yeast cells. eLife. 10, e62084 (2021).

57. J. Schindelin, I. Arganda-Carreras, E. Frise, V. Kaynig, M. Longair, T. Pietzsch, S. Preibisch, C. Rueden, S. Saalfeld, B. Schmid, J.-Y. Tinevez, D. J. White, V. Hartenstein, K. Eliceiri, P. Tomancak, A. Cardona, Fiji: an open-source platform for biological-image analysis. Nat Methods. 9, 676–682 (2012).

58. A. K. Lancaster, A. Nutter-Upham, S. Lindquist, O. D. King, PLAAC: a web and command-line application to identify proteins with prion-like amino acid composition. Bioinformatics. 30, 2501–2502 (2014).

