## Supplemental Text, Figures and Tables for "Force redistribution in clathrin-mediated endocytosis revealed by coiled-coil force sensors"

**This PDF file includes:**

Supplementary Text
Figs. S1 to S10
Table S1 to S2
Captions for Movies S1 to S3

**Other Supplementary Materials for this manuscript include the following:**

Movies S1 to S3

**Supplementary Text**

Calculation of the maximum unfolding force as a function of the force loading rate

The probability of unfolding of a given coiled-coil is governed by the following equation

$$\frac{dP_u}{dt} = k(t)[1 - P_u], \text{ where } k(t) = k_0 e^{\frac{F\Delta x}{k_B T}}$$

Changing the variable from $t$ to $F$ using $F = rt$, where $r$ is the loading rate and integrating the above equation, we get the probability of unfolding as a function of force as

$$P_u(F) = 1 - \exp\left[-\frac{k_0 k_B T}{\mathrm{r}\Delta x}\left(e^{\frac{F\Delta x}{k_B T}} - 1\right)\right]$$

By taking the derivative of $P_u(F)$ with respect to $F$, we can get the probability density as

$$PDF(F) = \frac{\mathrm{k}_0}{r}\exp\left[\frac{F\Delta x}{k_B T}\right]\exp\left[-\frac{k_0 k_B T}{\mathrm{r}\Delta x}\left(e^{\frac{F\Delta x}{k_B T}} - 1\right)\right]$$

The above function was used to fit the experimental PDF of unfolding forces (Fig. S10a-b). for cc-11pN. For cc-11pN, we can get $\Delta x$ =2.32 ± 0.48 nm and $k_0$= 4.35 ± 5.09 $10^{-3}$ $s^{-1}$.

Using the parameters extracted from fitting the experimental data, we can find the maximum unfolding force $F^*$ as a function of force loading rate $r$. $F^*$ is calculated from setting the first derivative of the PDF to zero, i.e.

$$\frac{d}{dF}[PDF(F)]_{F=F^*} = \frac{\mathrm{d}}{dF}\left[\frac{\mathrm{k}_0}{r}\exp\left[\frac{F\Delta x}{k_B T}\right]\exp\left[-\frac{k_0 k_B T}{\mathrm{r}\Delta x}\left(e^{\frac{F\Delta x}{k_B T}} - 1\right)\right]\right]_{F=F^*} = 0$$

Solving for $F^*$ readily gives

$$F^* = \frac{k_B T}{\Delta x}\ln\left[\frac{r\Delta x}{k_0 k_B T}\right]$$

For cc-11pN, the relationship between $F^*$ and force loading rate is shown in Fig. S10c. From the graph above, it can be seen that variations of the pulling rate in the physiological range (5-10 pN/s) changes $F^*$ by ~1 pN. This demonstrates that cc-11pN acts as a reliable force sensor for mechanosensitive proteins *in vivo*.

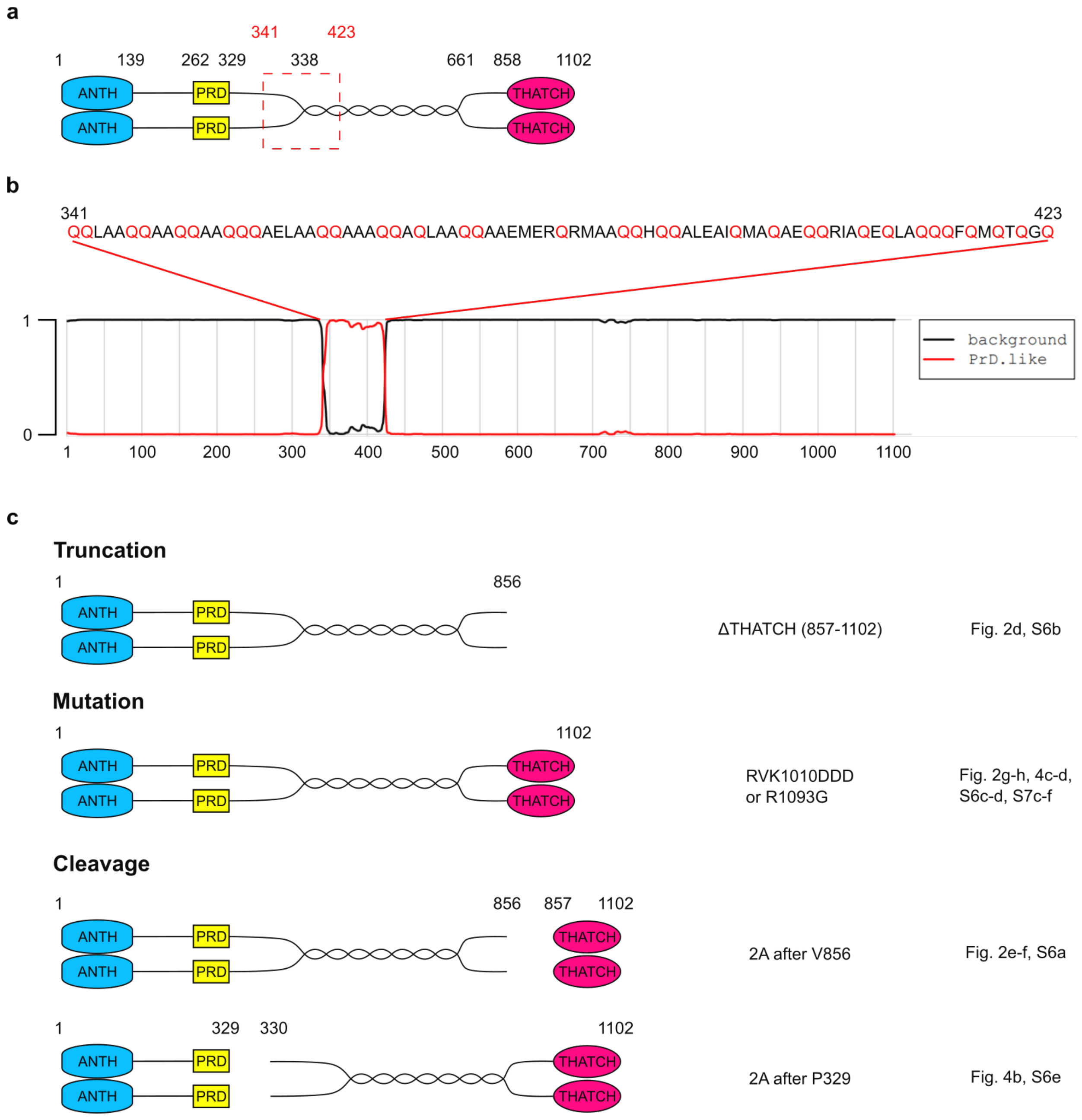


**Fig. S1. Domain organization of End4p.**

**a**, Schematic of End4p dimer. The numbers indicate the residues where domains start and end. The coiled-coil force sensors are inserted after D155, P337, or V856. The red dotted box denotes the poly-Q region in **b**. Drawings are not to scale. **b**, The beginning of End4p dimerization coiled-coil contains a poly-Q region that is predicted to be prion like by Prion Like Amino Acid Composition (PLAAC)(*58*). Glutamines (Q) are highlighted in red. **c**, The truncation, mutations and cleavages on End4p used in this study are shown to indicate the amino acid positions of these manipulations. Figure panel numbers that contain each manipulation are shown on the right.

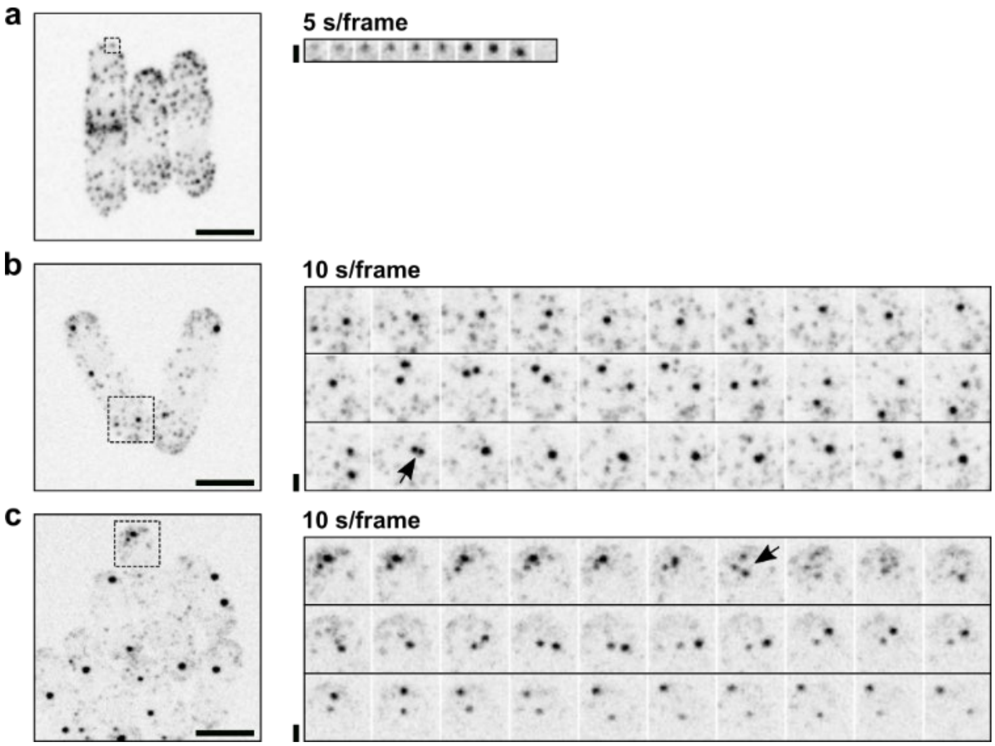


**Fig. S2. End4p constructs in condensates have longer lifetime than End4p at endocytic sites.**

**a**, Representative fission yeast cells with fluorescently tagged wild-type End4p. Right: temporal evolution of End4p in the boxed region. A typical End4p patch lasts less than 50 seconds. See also movie 1. **b**-**c**, Representative fission yeast cells with fluorescently-tagged End4p where a cc-11pN was inserted before the THATCH domain. Right: temporal evolution of End4p construct in the boxed regions. End4p condensates experience rapid movements across the cell and occasionally fuse with each other (**b**) or split (**c**) (arrows). A subset of this montage is shown in Fig. 1**g**. See also movie 2 (**b**) and movie 3 (**c**). **a-c**, scale bars in the left panels, 5 µm. Scale bars in the right panels, 2 µm. End4p is tagged at the C-terminal with mEGFP in **a**-**c**. Images are inverted contrast of max projected whole cell sections. Note that the contrast is enhanced in **a** to better visualize a single endocytic event.

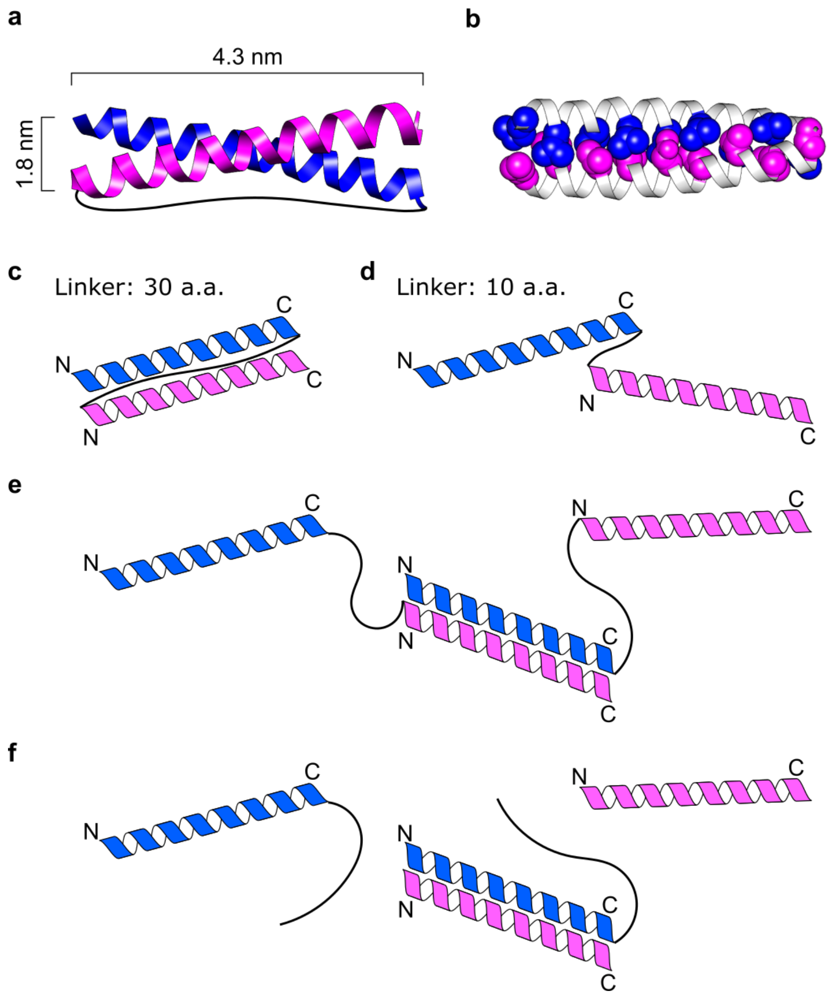


**Fig. S3. Schematic of coiled-coil force sensors in different conformations.**

**a**, Ribbon diagram of cc-11pN in the folded state. The two α-helices (blue and magenta) are connected with a 30-amino acid linker (black line). Dimensions of the coiled-coil are indicated according to the structure of PDB entry 2zta. **b**, Structure of a folded cc-11pN with the residues in the hydrophobic core highlighted. These hydrophobic residues are exposed when the coiled-coil unfolds under sufficiently large force. **c**, cc-11pN with a 30-amino acid linker folds properly by forming a parallel coiled-coil. **d**, cc-11pN with a 10-amino acid is unable to form a parallel coiled-coil and adopts an open conformation. This open conformation promotes the formation of protein condensates as shown in Fig. 2**a**-**b** and Fig. S4**h**. **e**, An intermolecular connection between two open cc-11pN is formed during the refolding process. Each intermolecular connection generates two exposed α-helices that could mediate further intermolecular connections. **f**, Cleavage at the linker of the two α-helices of cc-11pN prevents the formation of intermolecular connections. See also Fig. 2**f**. The N- and C-terminals of α-helices in **c**-**f** are labeled for orientation.

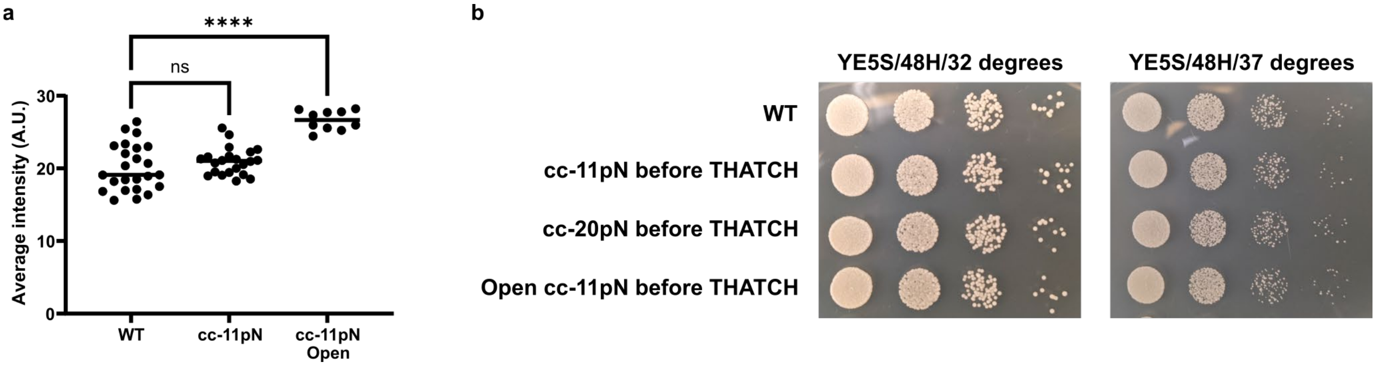


**Fig. S4. Comparison of End4p protein expression levels and cell growth in different strains.** **a**, The protein expression level of End4p was quantified by summing the whole cell fluorescence of C-terminally tagged End4p with mEGFP after background subtraction. WT: no insertion of coiled-coil force sensor (representative image in Fig. 1**f**). cc-11pN: 11pN coiled-coil force sensor inserted before THATCH (representative image in Fig. 1**g**). cc-11pN open: open conformation cc-11pN inserted before THATCH (representative image in Fig. 2**b**). Dunn's multiple comparison. ****: P<0.0001. Each dot represents one field of view. >200 cells for each group from at least three independent repeats. **b**, Growth assay of different strains was performed on YE5S plate for 48 hours at 32 degrees and 37 degrees. The growth rate of cells with unfolded coiled-coil (cc-11pN), folded coiled-coil (cc-20pN) or always open coiled-coil (open cc-11pN) were comparable with cells without the coiled-coil insertion. Data are representative of two independent repeats.

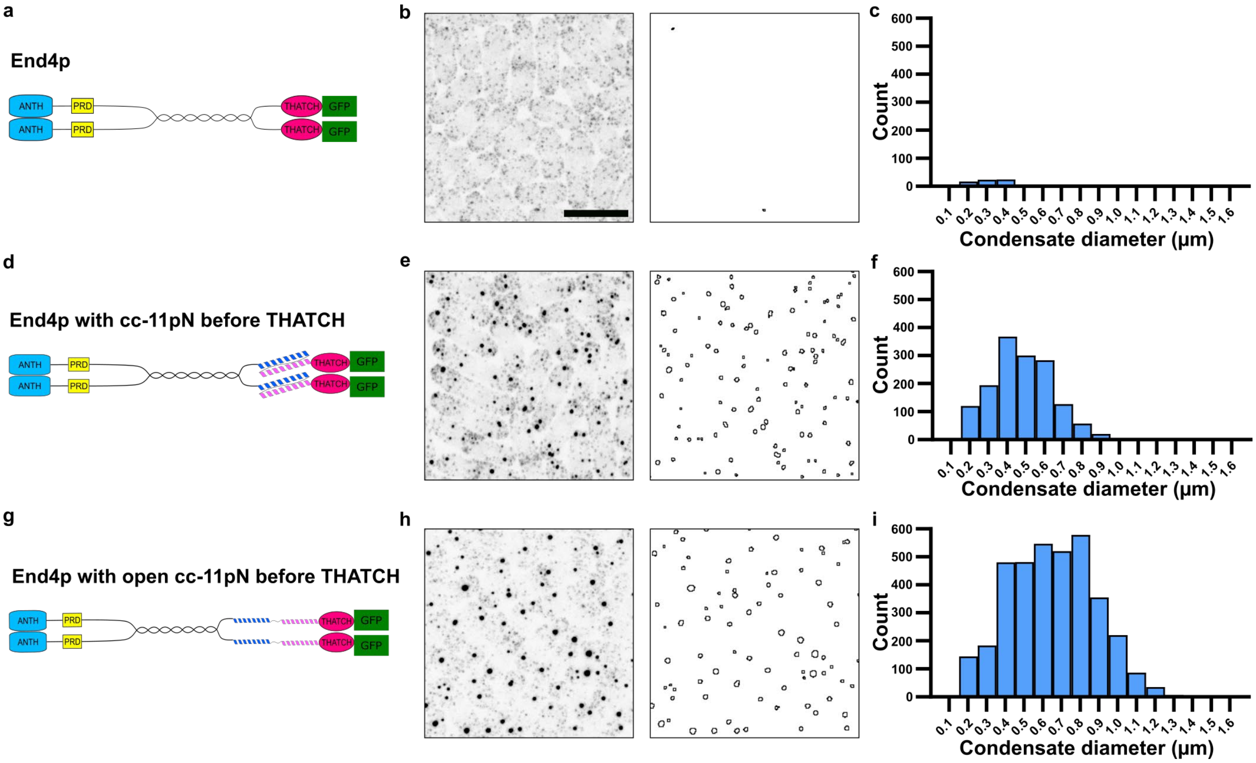


**Fig. S5. Quantification of protein condensates.**

End4p condensates were distinguished from End4p patches by comparing their size using a custom ImageJ plugin. **a-c,** End4p was tagged at the N-terminal with mEGFP. **d-f,** End4p was tagged at the N-terminal with mEGFP, and a cc-11pN was inserted before the THATCH domain. **g-i,** End4p was tagged at the N-terminal with mEGFP, and a cc-11pN in the open conformation was inserted before the THATCH domain. **b, e, h,** Representative images of fission yeast cells with fluorescently tagged End4p or End4p constructs are shown on the left, and the identified End4p condensates are shown on the right. The distributions of condensate diameter are shown in **c, f,** and **i**, each from >500 cells. A negligible population of End4p patches, typically from spatially overlapping endocytic events, is misidentified as End4p condensates in **b-c**. Scale bar in **b** also applies **e** and **h**, 10 µm. Images in **b, e, h** were acquired and displayed with the same settings. The same analysis was used for Fig. 3**b**.

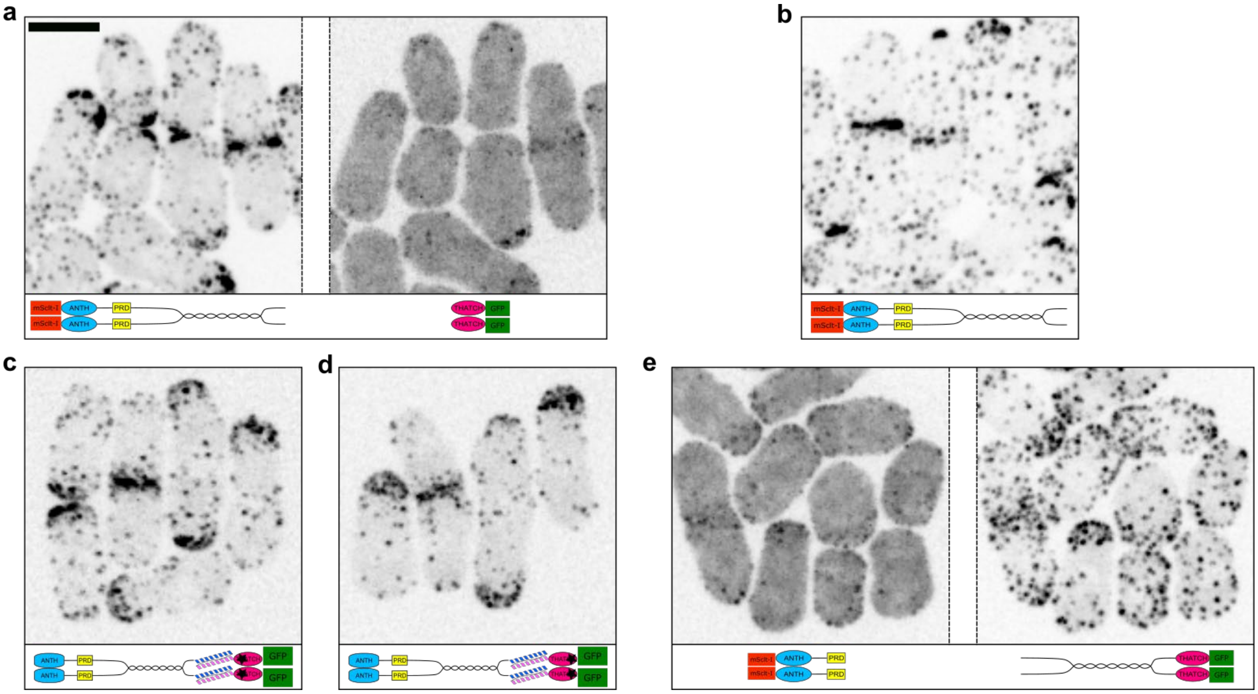


**Fig. S6. Controls for End4p localization in mutant strains.**

**a**, Representative fission yeast cells expressing End4p where a self-cleaving 2A peptide was inserted before the THATCH domain and fluorescently-tagged at its N- and C-terminals. The End4p N-terminal abnormally accumulates at cell tips and the division plane, while End4p C-terminal is diffusive in the cytoplasm. The localization of End4p fragments were not changed by the insertion of cc-11pN as shown in Figure 2**e**. **b**, Representative fission yeast cells expressing fluorescently-tagged End4p where the THATCH domain was deleted. End4p without the THATCH domain abnormally accumulates at cell tips and the division plane. The localization of this End4p construct was not changed by the insertion of cc-11pN as shown in Figure 2**d**. **c**, Representative fission yeast cells expressing fluorescently-tagged End4p where the RVK1010DDD mutation was introduced into the THATCH domain to abolish binding to actin filaments. This mutant abnormally accumulates at cell tips and the division plane. **d**, Representative fission yeast cells expressing fluorescently-tagged End4p where the R1093G mutation was introduced into the THATCH domain to abolish binding to actin filaments. This mutant abnormally accumulates at cell tips and the division plane. The localization of End4p in **c** and **d** were not changed by the insertion of cc-11pN as shown in Figure 2**g** and 2**h**. **e**, Representative fission yeast cells expressing fluorescently-tagged End4p where a self-cleaving 2A peptide was inserted after the proline rich domain. End4p N-terminal is diffusive in the cytoplasm, while its C-terminal localizes to endocytic patches. Compare with Figure 4**b**. **a-e**, Scale bar in **a** applies to all images, 5 µm. Images are inverted contrast of max projected whole cell z-stacks. End4p is tagged at the N-terminal with mScarlet-I in **a, b,** and **e**, and End4p is tagged at the C-terminal with mEGFP in **a, c, d** and **e**. The schematic under each image indicates the modifications on End4p.

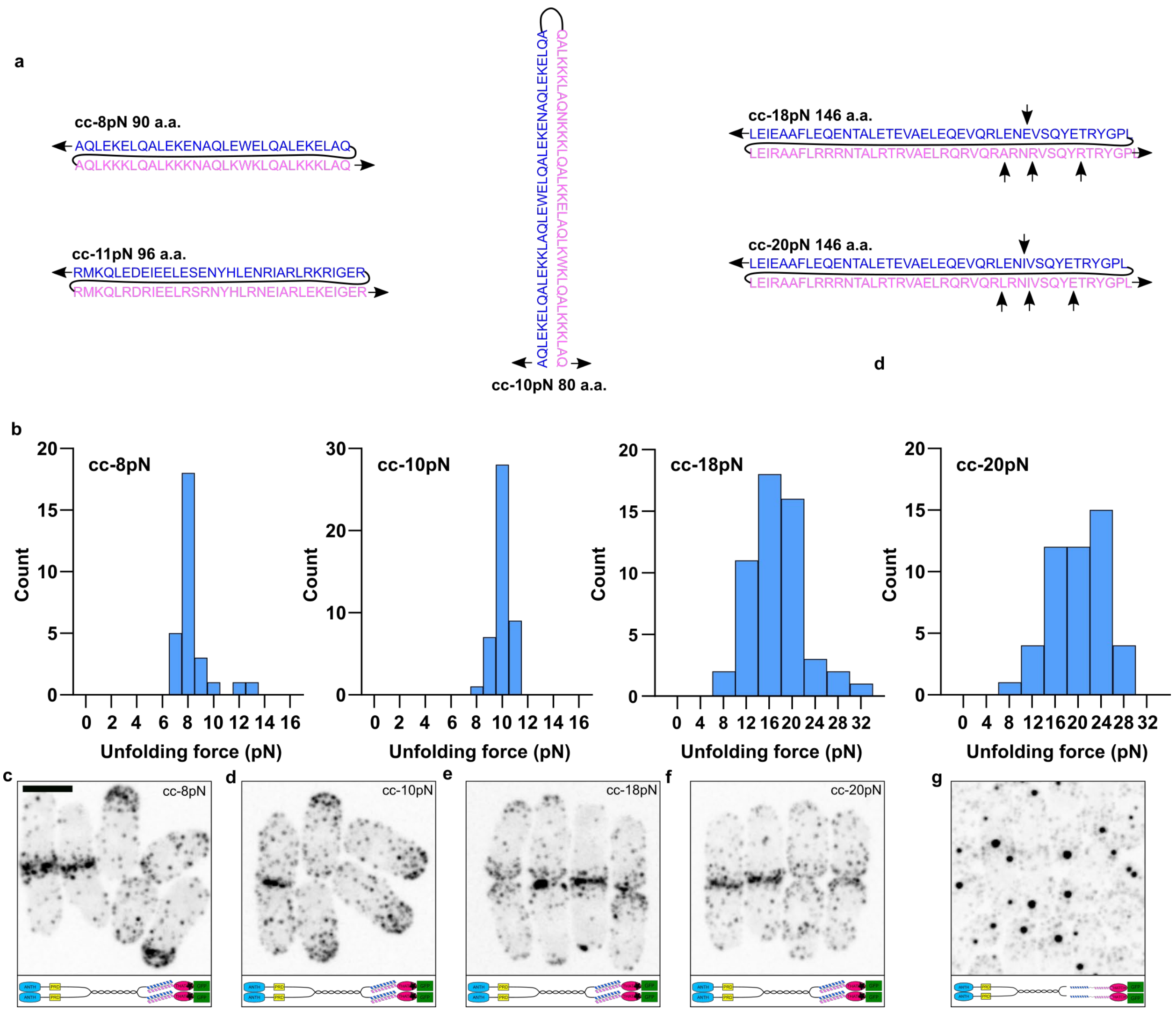


**Fig. S7. Coiled-coils with different average unfolding forces.**

**a,** Sequence and pulling direction (indicated by black arrows) of coiled-coil force sensors. Among the five force sensors, cc-8pN, cc-11pN, cc-18pN and cc-20pN are parallel coiled-coils while cc-10pN is an anti-parallel coiled-coil. The flexible linker between the two α-helices (blue and magenta) are drawn as black lines. Amino acids that were mutated between cc-18pN and cc-20pN are labeled by vertical arrows. **b**. Distribution of the unfolding force for cc-8pN, cc-10pN, cc-18pN and cc-20pN. Data were used to calculate the average unfolding force as shown in Figure 3**a**. **c-f**, Representative fission yeast cells expressing fluorescently-tagged End4p where cc-8 pN, cc-10 pN, cc-18pN or cc-20 pN were inserted before the THATCH domain, and an R1093G mutation was introduced into the THATCH domain to abolish its binding to actin filaments. End4p shows abnormal accumulation at cell tips and the division plane, without forming condensates. **g**. Representative fission yeast cells expressing fluorescently-tagged End4p where a variant of cc-20pN that is always in an open conformation was inserted before the THATCH domain. The insertion of this "always-open" coiled-coil into End4p led to End4p condensates. **c-g**, Scale bar in **c** applies to all images, 5 μm. Images are inverted contrast of maximm projected whole cell sections. The schematic under each image indicates the modifications on End4p.

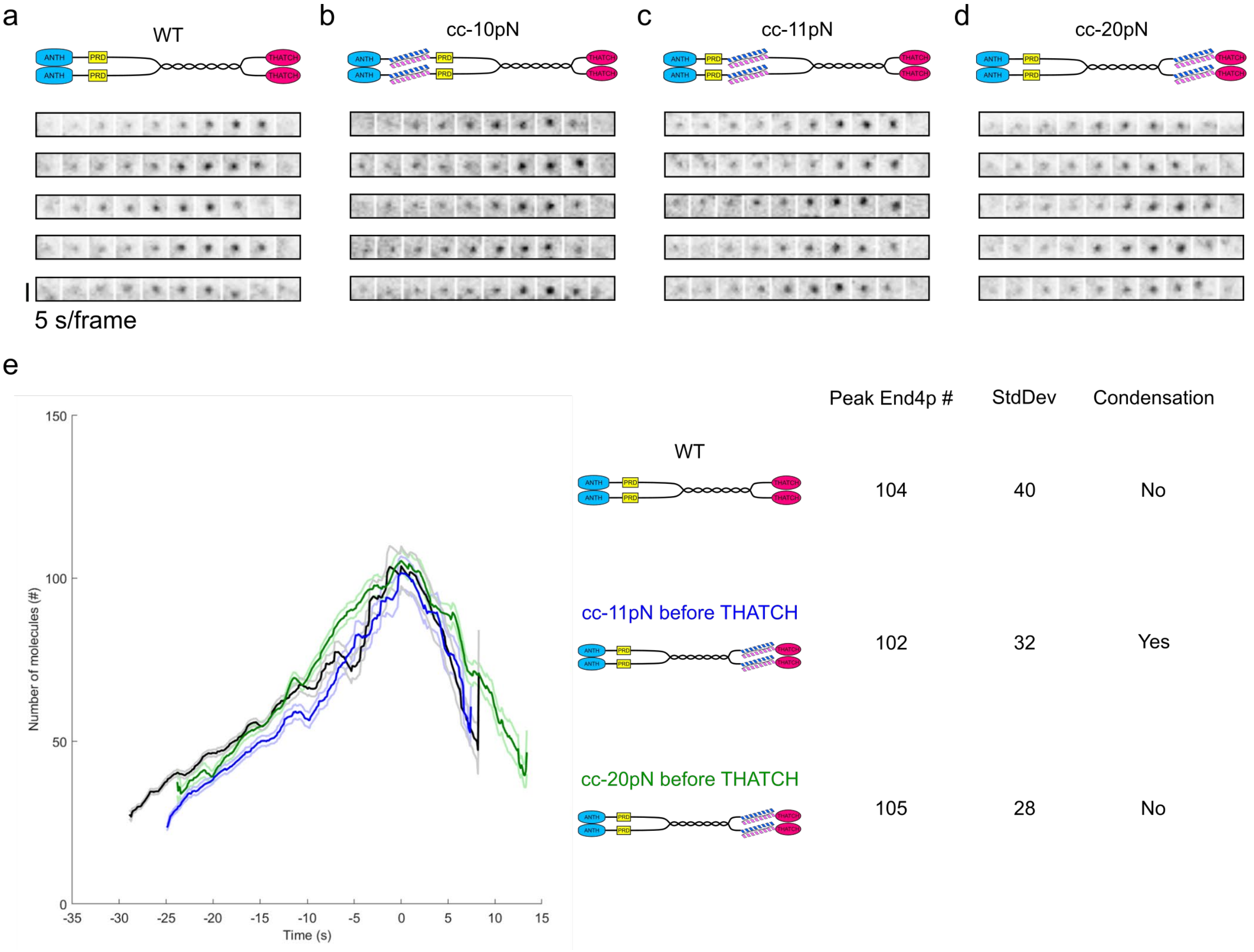


**Fig. S8. Temporal evolution of End4p patches at endocytic sites.**

**a-d,** The temporal evolution of End4p with or without coiled-coil force sensors is recorded at endocytic sites. The insertion of a coiled-coil force sensor with an unfolding force threshold higher than the local force does not affect the assembly and disassembly of End4p molecules. Scale bar, 2 µm. End4p is tagged at the C-terminal with mEGFP in **a-d**. Images are inverted contrast of maximum projection of z-stacks covering the whole cell. The schematic above each image indicates the location of coiled-coil force sensors in End4p, and fluorescent tags are omitted for clarity. **e,** Temporal evolution of the number of End4p molecules at endocytic patches determined by patch tracking (*55*). There is no difference between the different constructs. Black: wild-type End4p (N=168; wild-type End4p does not form condensates), blue: End4p with cc-11pN inserted before THATCH (N=100; this construct form condensates), green: End4p  with cc-20pN inserted before THATCH (N=104; note this construct does not form condensates). Quantifications of peak End4p number are listed at the right of the curve. Dark colors correspond to averages and light colors correspond to confidence intervals at 95%. Welch's t-test was used to compare the mean peak number of molecules. Time 0 s corresponds to the peak of End4p.

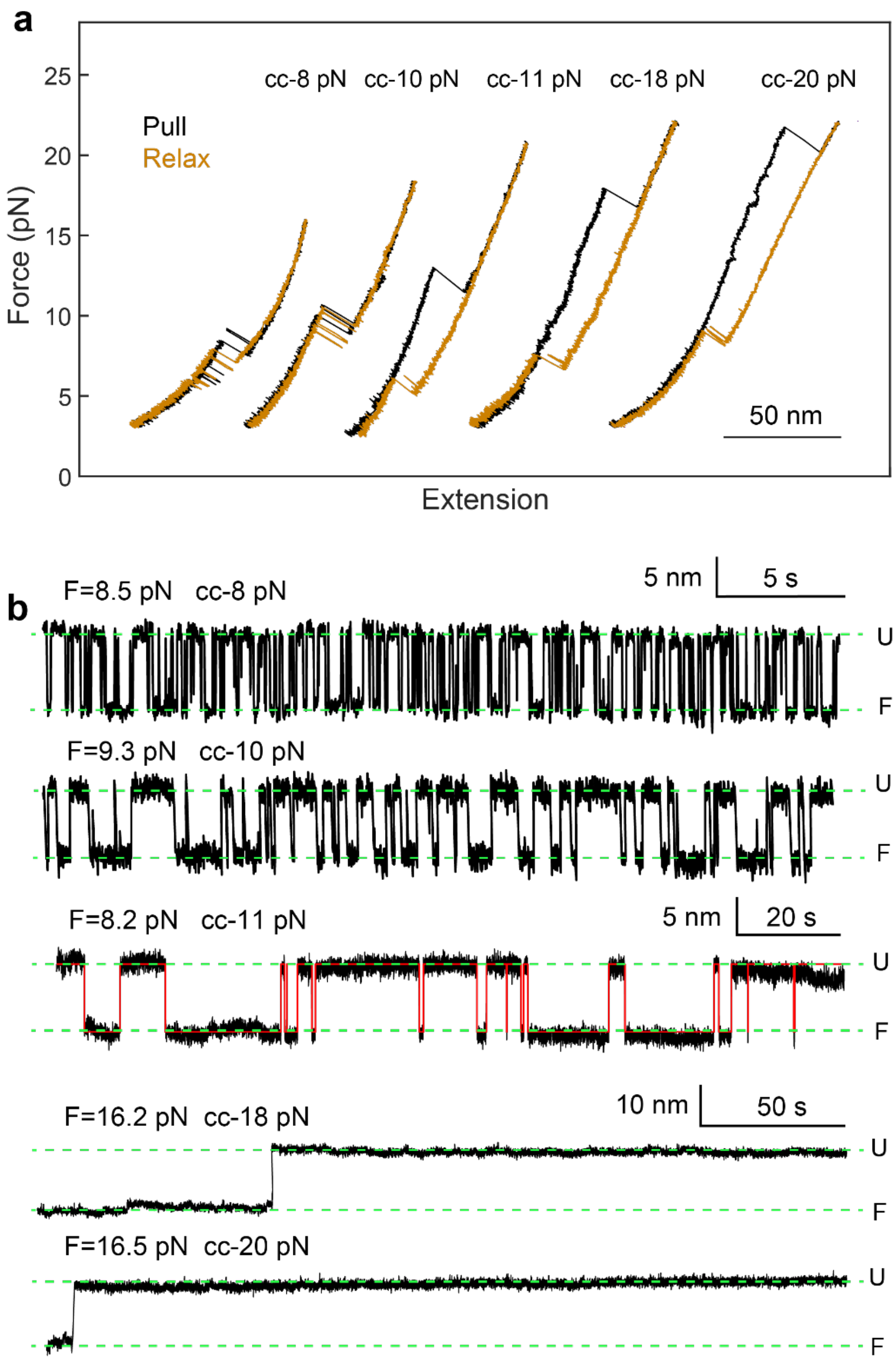


**Fig. S9. Unfolding and refolding transitions of coiled coils in or out of thermal equilibrium.** **a,** Force-extension curves of the coiled-coils showing minimum (for cc-8pN and cc-10pN) or pronounced (for cc-11pN, cc-18pN and cc-20pN) hysteresis between pulling (black) and relaxation (orange) curves. **b,** Time-dependent extension transitions of the coiled-coils at constant mean force showing folding and unfolding transitions under thermal equilibrium for cc-8pN, cc-10pN, and cc-11pN or out of thermal equilibrium for cc-18pN and cc-20pN. The equilibrium trajectories were analyzed by hidden-Markov modeling, leading to the idealized transition shown in red lines and the corresponding state lifetimes (Table S2). The green dashed lines mark the average positions of the folded states and unfolded states, indicated by “F” and “U”, respectively, on the right.

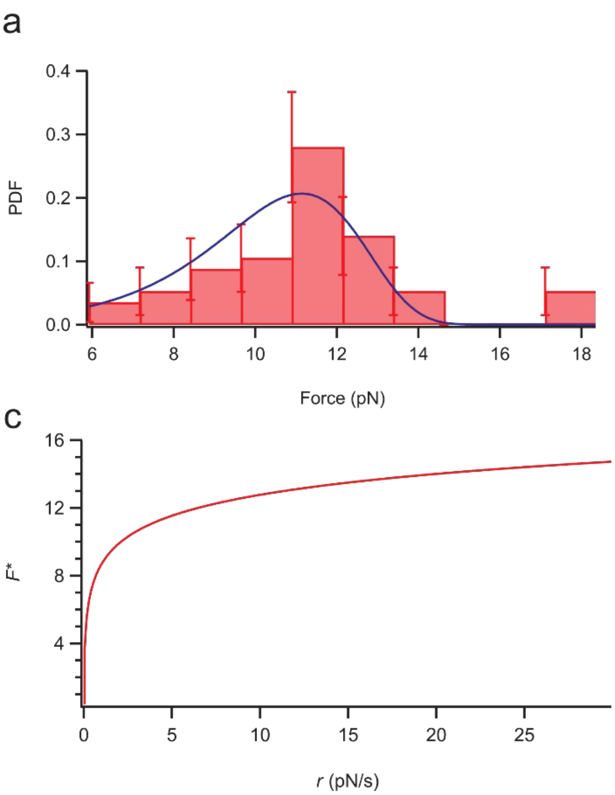


**Fig. S10. The unfolding force of a coiled-coil is insensitive to pulling rate above 5pN/s. a,** Fitting of PDF (blue line) to experimental data of the unfolding force of cc-11pN. **b,** Probability of coiled-coil opening as a function of the pulling force. **c,** Relationship between the maximum unfolding force and the pulling rate. F* is insensitive to pulling rate >5pN/s.

**Table S1.** *S. pombe* strains used in this study.

| Strain | Genotype | Reference |
|---|---|---|
| JB367 | end4-mEGFP fex1Δ fex2Δ ade6-M216 his3-D1 leu1-32 ura4-D18 h- | This study |
| JB561 | mScarlet-I-end4-mEGFP fex1Δ fex2Δ ade6-M216 his3-D1 leu1-32 ura4-D18 h- | This study |
| JB562 | mScarlet-I-end4 fex1Δ fex2Δ ade6-M216 his3-D1 leu1-32 ura4-D18 h- | This study |
| JB577 | mScarlet-I-end4-ERBV-1-20AA-end4-mEGFP fex1Δ fex2Δ ade6-M216 his3-D1 leu1-32 ura4-D18 h- | This study |
| JB586 | mScarlet-I-end4ΔC fex1Δ fex2Δ ade6-M216 his3-D1 leu1-32 ura4-D18 h- | This study |
| JB592 | mScarlet-I-end4-GCN4-pIL-I16N-5E3R-2A-GCN4-pIL-I16N-5R3E-end4-mEGFP fex1Δ fex2Δ ade6-M216 his3-D1 leu1-32 ura4-D18 h- | This study |
| JB597 | mScarlet-I-end4-GCN4-pIL-I16N-5E3R-2A-MUT-GCN4-pIL-I16N-5R3E-end4-mEGFP fex1Δ fex2Δ ade6-M216 his3-D1 leu1-32 ura4-D18 h- | This study |
| JB617 | mScarlet-I-end4-Oakley-p1-OnePlus-end4-mEGFP fex1Δ fex2Δ ade6-M216 his3-D1 leu1-32 ura4-D18 h- | This study |
| JB619 | mScarlet-I-end4-GCN4-pIL-I16N-5E3R-10G-GCN4-pIL-I16N-5R3E-end4-mEGFP fex1Δ fex2Δ ade6-M216 his3-D1 leu1-32 ura4-D18 h- | This study |
| JB643 | end4-Velcro-p1-end4-mEGFP fex1Δ fex2Δ ade6-M216 his3-D1 leu1-32 ura4-D18 h- | This study |
| JB675 | mScarlet-I-end4-GCN4-pIL-I16N-5E3R-2A_Mut-pIL-I16N-5R3E fex1Δ fex2Δ ade6-M216 his3-D1 leu1-32 ura4-D18 h- | This study |
| JB689 | mScarlet-I-end4-ERBV1-20AA-GCN4-pIL-I16N-5E3R-2A_Mut-GCN4-pIL-I16N-5R3E-end4-mEGFP fex1Δ fex2Δ ade6-M216 his3-D1 leu1-32 ura4-D18 h- | This study |
| JB691 | mScarlet-I-end4-ANTH-GCN4-pIL-I16N-5E3R-2A_Mut-GCN4-pIL-I16N-5R3E-end4-mEGFP fex1Δ fex2Δ ade6-M216 his3-D1 leu1-32 ura4-D18 h- | This study |
| JB693 | mScarlet-I-end4-ANTH-Pro-ERBV1-20AA-end4-mEGFP fex1Δ fex2Δ ade6-M216 his3-D1 leu1-32 ura4-D18 h- | This study |
| JB694 | mScarlet-I-end4-ANTH-Pro-GCN4-pIL-I16N-5E3R-2A_Mut-GCN4-pIL-I16N-5R3E-end4-mEGFP fex1Δ fex2Δ ade6-M216 his3-D1 leu1-32 ura4-D18 h- | This study |
| JB717 | mScarlet-I-end4-pER-E-GGGS-pER-R-end4-mEGFP fex1Δ fex2Δ ade6-M216 his3-D1 leu1-32 ura4-D18 h- | This study |
| JB726 | mScarlet-I-end4-1010DDD-mEGFP fex1Δ fex2Δ ade6-M216 his3-D1 leu1-32 ura4-D18 h- | This study |
| JB727 | mScarlet-I-end4-V856-GCN4-pIL-I16N-5E3R-2A_Mut-GCN4-pIL-I16N-5R3E-1010DDD-mEGFP fex1Δ fex2Δ ade6-M216 his3-D1 leu1-32 ura4-D18 h- | This study |
| JB728 | mScarlet-I-end4-R1093G-mEGFP fex1Δ fex2Δ ade6-M216 his3-D1 leu1-32 ura4-D18 h- | This study |

| | | |
|---|---|---|
| JB729 | mScarlet-I-end4-V856-GCN4-pIL-I16N-5E3R-2A_Mut-GCN4-pIL-I16N-5R3E-R1093G-mEGFP fex1Δ fex2Δ ade6-M216 his3-D1 leu1-32 ura4-D18 h- | This study |
| JB730 | mScarlet-I-end4-GCN4-pIL-I16N-5E3R-2A_Mut-GCN4-pIL-5R3E-ΔC fex1Δ fex2Δ ade6-M216 his3-D1 leu1-32 ura4-D18 h- | This study |
| JB735 | mScarlet-I-end4-pER-E-10G-pER-R-end4-mEGFP fex1Δ fex2Δ ade6-M216 his3-D1 leu1-32 ura4-D18 h- | This study |
| JB736 | mScarlet-I-end4-ANTH-Pro-ERBV1-20AA-V856-GCN4-pIL-I16N-5E3R-2A_Mut-GCN4-pIL-I16N-5R3E-end4-mEGFP fex1Δ fex2Δ ade6-M216 his3-D1 leu1-32 ura4-D18 h- | This study |
| JB743 | mScarlet-I-end4-ANTH-Pro-Velcro-p1-end4-mEGFP fex1Δ fex2Δ ade6-M216 his3-D1 leu1-32 ura4-D18 h- | This study |
| JB753 | mScarlet-I-end4-ANTH-Pro-Velcro-p1-end4-R1093G-mEGFP fex1Δ fex2Δ ade6-M216 his3-D1 leu1-32 ura4-D18 h- | This study |
| JB760 | mScarlet-I-end4-ANTH-Velcro-p1-end4-mEGFP fex1Δ fex2Δ ade6-M216 his3-D1 leu1-32 ura4-D18 h- | This study |
| JB762 | mScarlet-I-end4-ANTH-Pro-Oakley-p1-OnePlus-end4-mEGFP fex1Δ fex2Δ ade6-M216 his3-D1 leu1-32 ura4-D18 h- | This study |
| JB767 | mScarlet-I-end4-ANTH-Oakley-p1-OnePlus-end4-mEGFP fex1Δ fex2Δ ade6-M216 his3-D1 leu1-32 ura4-D18 h- | This study |
| JB772 | mScarlet-I-end4-ANTH-pER-E-GGGS-pER-R-end4-mEGFP fex1Δ fex2Δ ade6-M216 his3-D1 leu1-32 ura4-D18 h- | This study |
| JB773 | mScarlet-I-end4-ANTH-Pro-pER-E-GGGS-pER-R-end4-mEGFP fex1Δ fex2Δ ade6-M216 his3-D1 leu1-32 ura4-D18 h- | This study |
| JB780 | mScarlet-I-end4-ANTH-Velcro-p1-R1093G-end4-mEGFP fex1Δ fex2Δ ade6-M216 his3-D1 leu1-32 ura4-D18 h- | This study |
| JB798 | mScarlet-I-end4-Oakley-p1-OnePlus-R1093G-end4-mEGFP fex1Δ fex2Δ ade6-M216 his3-D1 leu1-32 ura4-D18 h- | This study |
| JB799 | mScarlet-I-end4-Velcro-p1-R1093G-end4-mEGFP fex1Δ fex2Δ ade6-M216 his3-D1 leu1-32 ura4-D18 h- | This study |
| JB800 | mScarlet-I-end4-pER-E-GGGS-pER-R-R1093G-end4-mEGFP fex1Δ fex2Δ ade6-M216 his3-D1 leu1-32 ura4-D18 h- | This study |
| JB805 | ent1-P571-STOP mEGFP-end4-pER-E-GGGS-pER-R-end4-mEGFP fex1Δ fex2Δ ade6-M216 his3-D1 leu1-32 ura4-D18 | This study |
| JB833 | ent1-P571-STOP mScarlet-I-end4-ANTH-Oakley-p1-OnePlus-end4-mEGFP fex1Δ fex2Δ ade6-M216 his3-D1 leu1-32 ura4-D18 | This study |
| JB834 | ent1-P571-STOP mScarlet-I-end4-ANTH-Pro-GCN4-pIL-I16N-5E3R-2A_Mut-GCN4-pIL-I16N-5R3E-end4-mEGFP fex1Δ fex2Δ ade6-M216 his3-D1 leu1-32 ura4-D18 | This study |
| JB848 | mScarlet-I-end4-pER-E-GGGS_pER-R-LA-end4-mEGFP fex1Δ fex2Δ ade6-M216 his3-D1 leu1-32 ura4-D18 | This study |
| JB1087 | end4-V856-mTq2-HP35-mNG-end4 fex1Δ fex2Δ ade6-M216 his3-D1 leu1-32 ura4-D18 | This study |
| JB1088 | end4-V856-mTq2-HP35st-mNG-end4 fex1Δ fex2Δ ade6-M216 his3-D1 leu1-32 ura4-D18 | This study |

| | | |
|---|---|---|
| JB1102 | pil1::mTq2-HP35-mNG fex1Δ fex2Δ ade6-M216 his3-D1 leu1-32 ura4-D18 | This study |
| JB1103 | pil1::mTq2-HP35st-mNG fex1Δ fex2Δ ade6-M216 his3-D1 leu1-32 ura4-D18 | This study |
| JB1116 | end4-V856-mTq2-end4 fex1Δ fex2Δ ade6-M216 his3-D1 leu1-32 ura4-D18 | This study |
| JB1117 | pil1::mTq2 fex1Δ fex2Δ ade6-M216 his3-D1 leu1-32 ura4-D18 | This study |
| JB1153 | mScarlet-I-end4-pER-E-GGGS_pER-R-LA-end4-R1093G-mEGFP fex1Δ fex2Δ ade6-M216 his3-D1 leu1-32 ura4-D18 | This study |

**Table S2.** Equilibrium forces and lifetimes of the calibrated coiled-coil force sensors. N.A., not measured due to the extremely slow transitions at the equilibrium force (see Fig. S9). Note that the force derived from the formation of End4p condensates in vivo correlates better with the unfolding force of the coiled coil sensors (indicated in the name of the sensor), instead of their equilibrium force.

| **Force sensors** | cc-8 pN | cc-10 pN | cc-11 pN | cc-18 pN | cc-20 pN |
|---|---|---|---|---|---|
| Equilibrium force (pN) | 8.4 (0.1) | 9.8 (0.2) | 8.4 (0.2) | N.A. | N.A. |
| State lifetime (s) | 0.09 | 0.45 | 9.1 | N.A. | N.A. |

**Movie S1.**

Time lapse of fission yeast cells where End4p is tagged with mEGFP. Images are inverted contrast of max projected whole cell sections. Scale bar, 5 μm. Time stamp, minute: second.

**Movie S2.**

Time lapse of fission yeast cells where End4p is tagged with mEGFP, and a cc-11pN was inserted before the THATCH domain. Images are inverted contrast of max projected whole cell sections. Scale bar, 5 μm. Time stamp, minute: second.

**Movie S3.**

Time lapse of fission yeast cells where End4p is tagged with mEGFP, and a cc-11pN was inserted before the THATCH domain. Images are inverted contrast of max projected whole cell sections. Scale bar, 5 μm. Time stamp, minute: second.
